# Structural mechanism of leaflet-specific phospholipid modulation of a pentameric ligand-gated ion channel

**DOI:** 10.1101/2022.06.07.494883

**Authors:** John T. Petroff, Noah M. Deitzen, Ezry Santiago-McRae, Brett Deng, Maya S. Washington, Lawrence J. Chen, K. Trent Moreland, Zengqin Deng, Michael Rau, James A. J. Fitzpatrick, Peng Yuan, Thomas T. Joseph, Jérôme Hénin, Grace Brannigan, Wayland W. L. Cheng

## Abstract

Pentameric ligand-gated ion channels (pLGICs) mediate synaptic transmission and are sensitive to their lipid environment. The mechanism of phospholipid modulation of any pLGIC is not well understood. We demonstrate that the model pLGIC, ELIC (*Erwinia* ligand-gated ion channel), is positively modulated by the anionic phospholipid, phosphatidylglycerol, from the outer leaflet of the membrane. To elucidate the mechanism of phosphatidylglycerol modulation, we determine a structure of ELIC in an open conformation. The structure shows a bound phospholipid in an outer leaflet site, and conformational changes in the phospholipid binding site unique to the open state. In combination with streamlined alchemical free energy perturbation calculations and functional measurements in asymmetric liposomes, the data support a mechanism by which an anionic phospholipid stabilizes the open state of a pLGIC by specific, state-dependent binding to this site.

## INTRODUCTION

pLGICs such as the nicotinic acetylcholine receptor (nAChRs) are ubiquitous in the nervous system where they determine synaptic transmission and neuronal excitability^1^. These channels are targets of general anesthetics and anti-epileptics^2^, and potential targets for the treatment of many neuropsychiatric diseases^3–5^. pLGIC function is modulated by certain lipids^6^, including anionic phospholipids which enhance agonist response in the nAChR^7–11^. Anionic phospholipids, as well as sterols and fatty acids, have also been shown to modulate other pLGICs^12–16^. It is intuitive that lipids modulate pLGICs by influencing the stability of different states that the channel samples during gating; however, mechanisms of phospholipid modulation of any pLGIC remain poorly understood.

With the recent boon in lipid-bound pLGIC structures by cryo-EM, investigation into the structural mechanisms of lipid modulation has focused on direct binding of lipids to specific sites^12, 13, 15, 17, 18^. Structures of the 5-HT_3A_ receptor (5-HT_3A_R), GABA_A_ receptor, and the prototypic pLGIC, *Erwinia chrysanthemi* ligand-gated ion channel (ELIC), show bound phospholipids at various sites^14, 19–23^. However, a phospholipid density in a structure is insufficient evidence for a modulatory mechanism^6^. Establishing such a mechanism demands knowledge of the functional effect of the phospholipid, the identity of the phospholipid that occupies the site, and state-dependent binding at that site consistent with the modulatory effect of the phospholipid. Recent structural studies of ELIC show a phospholipid and cardiolipin density bound to different sites in the apo or resting structure, but functional analysis indicates that cardiolipin stabilizes the open state of the channel^14^. Another study of the 5-HT_3A_R shows evidence of state-dependent phospholipid binding, but the identity of this phospholipid in the structure and the modulatory effect of phospholipids in 5-HT_3A_R is unknown^19^.

In this study, we determine a structural mechanism for anionic phospholipid positive modulation of the pLGIC, ELIC. An open structure of ELIC shows a phospholipid density in an outer leaflet M4-M3-M1 site not well resolved in non-conducting structures. The phospholipid binding site shows a distinct conformation in the open structure. Along with streamlined free energy perturbation calculations^24^ and functional measurements in asymmetric proteoliposomes the results support a mechanism by which the anionic phospholipid, phosphatidylglycerol (PG), stabilizes the open state of ELIC by binding to a specific site.

## Results

### Modulation of ELIC by neutral and anionic phospholipids

pLGICs such as the nAchR require an anionic phospholipid and phosphatidylethanolamine (PE) for maximal agonist response^7^. Using a fluorescence stopped-flow liposomal flux assay, we previously showed that ELIC agonist responses are significantly enhanced in 2:1:1 POPC (1-palmitoyl-2-oleoyl-phosphatidylcholine): POPE (1-palmitoyl-2-oleoyl-phosphatidylethanolamine): POPG (1-palmitoyl-2-oleoyl-phosphatidylglycerol) liposomes compared to POPC-only liposomes^13, 25^. This effect manifested as an increase in peak response and slowing of the decay in channel activity (Fig. 1a, 1c and 1d). However, the exact contributions of POPG and POPE on these effects were not clear.

**Fig. 1.**
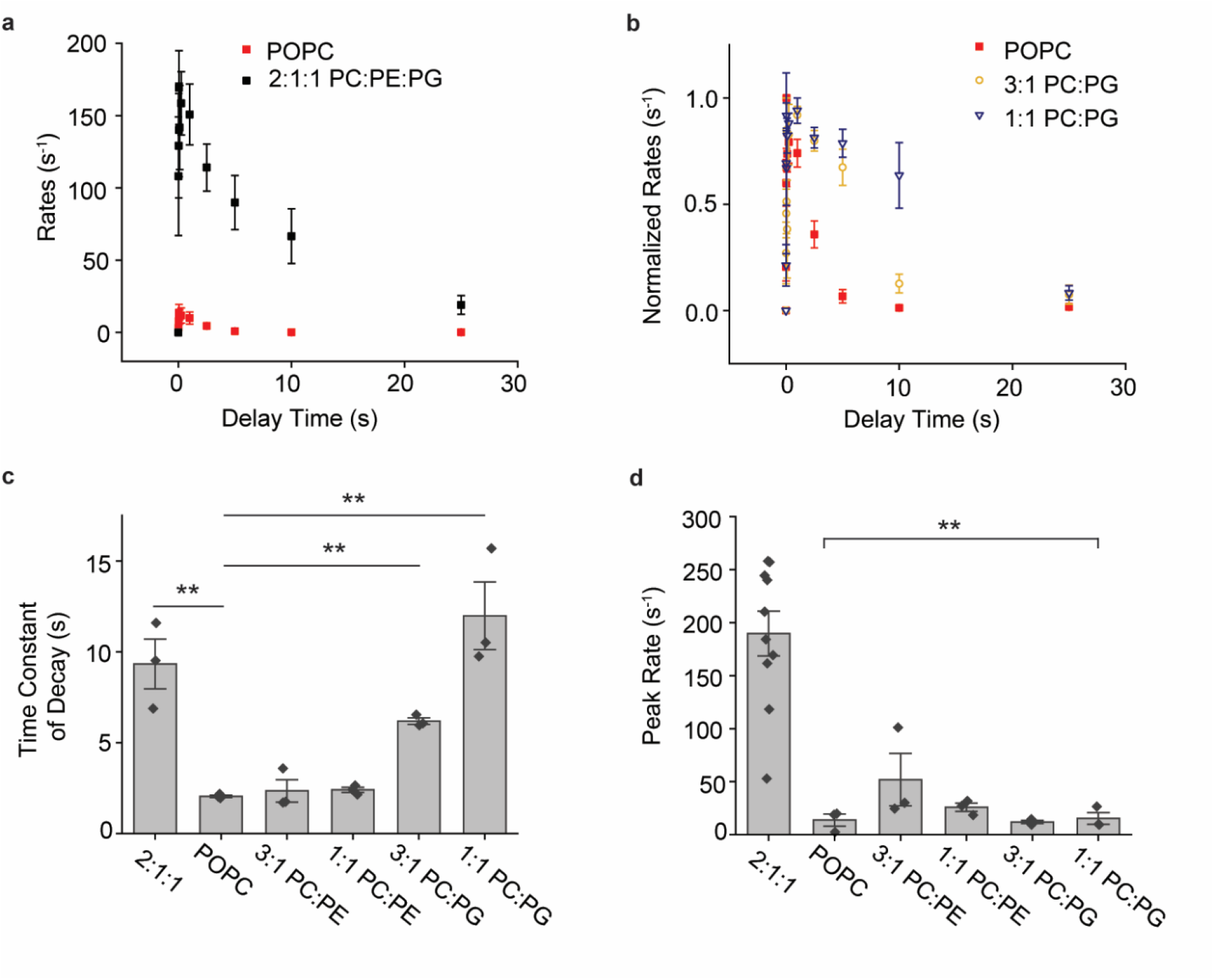
ELIC agonist responses in different lipid conditions. **(a)** Tl^+^ flux rates of WT ELIC in 2:1:1 POPC:POPE:POPG (black) and POPC (red) liposomes as a function of time after mixing with 10 mM propylamine (n = 3, ± SEM). **(b)** Normalized Tl^+^ flux rates of WT ELIC in POPC, 3:1 POPC:POPG and 1:1 POPC:POPG as a function of time after mixing with 10 mM propylamine (n = 3, ± SEM). **(c)** Weighted time constant of decay of channel activity (decay time courses were fit with a double exponential) in response to 10 mM propylamine from the Tl^+^ flux assay for the indicated liposome compositions (n = 3, ± SEM). 2:1:1 indicates 2:1:1 POPC:POPE:POPG. ** indicates p<0.01. **(d)** Same as (C) showing peak channel activity (rate) in response to 10 mM propylamine from the Tl+ flux assay (2:1:1, n = 12; other, n = 3, ± SEM). ** indicates p<0.01 when comparing 2:1:1 with all other liposome compositions.

To delineate the respective effects of these lipids, ELIC activity was measured in binary mixtures of 3:1 or 1:1 of POPC:POPE or POPC:POPG. Responses were measured using 10 mM propylamine (a saturating concentration of agonist, Extended Data Fig. 1a), since cysteamine (the agonist used previously^13^) at concentrations >5 mM is incompatible with thallium. Only POPG (25-50 mol% POPG) produced significant slowing of the decay in channel activity (Fig. 1b and 1c). Accordingly, POPG significantly increased relative channel activity at 10 and 25 s after agonist application, prolonging channel open times (Extended Data Fig. 1b-1c). Importantly, both POPG and POPE were required to achieve maximal responses (Fig. 1d). Furthermore, the EC_50_ for the peak agonist dose-response in POPC-only liposomes was ∼2x higher than in 2:1:1 POPC:POPE:POPG liposomes but not significantly different (Extended Data Fig. 1a), suggesting that POPE and POPG do not significantly shift the equilibrium between resting and agonist-bound states. Taken together, the results show that POPG modulates ELIC function by stabilizing the open state of the channel relative to one or more agonist-bound non-conducting states. This non-conducting state could be a desensitized state^13^ since POPG slows the rate of the decay in channel activity, or a pre-active state^12, 26^ since POPG enhances peak responses in the presence of POPE. While differences in channel reconstitution and orientation in the various liposome compositions could contribute to the effect on peak response from these experiments, experiments using asymmetric liposomes described below verify that POPG increases peak channel activity.

### Capturing an open conformation of ELIC

To investigate the mechanism by which POPG stabilizes the open state of ELIC, we determined structures of WT ELIC by single particle cryo-electron microscopy (cryo-EM) in the absence and presence of agonist in MSP1E3D1 nanodiscs with 2:1:1 POPC:POPE:POPG (hereafter referred to as 2:1:1) or POPC (Extended Data Fig. 2, Table 1). For all structures, 10 mM cysteamine was used as the agonist since this is a saturating concentration of the most potent, known agonist of ELIC^27^. The POPC and 2:1:1 WT ELIC apo structures were indistinguishable. Likewise, the agonist-bound structures were identical in both lipid conditions (Extended Data Fig. 3a), deviating insignificantly from apo and propylamine-bound structures in POPC nanodiscs previously reported^14, 28^. Importantly, the agonist-bound structures showed a non-conducting pore with occlusion at 16’ (F247) and 9’ (L240) (Fig. 2e and 2f). Thus, structures of WT ELIC in poorly activating POPC compared to activating 2:1:1 lipids exhibited no meaningful structural differences.

**Fig. 2.**
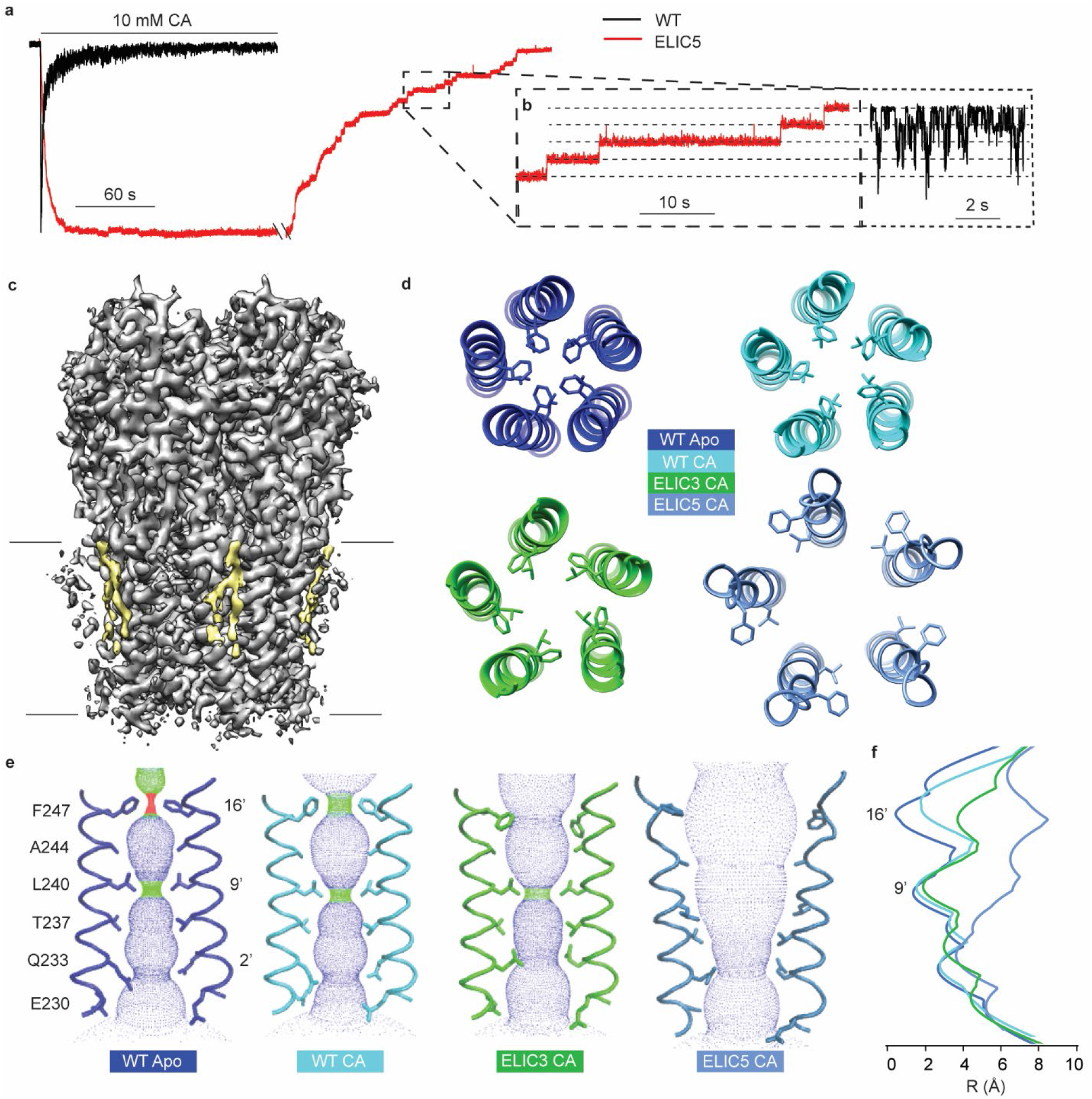
Conformational changes of pore opening. **(a)** Representative ELIC currents in response to 10 mM cysteamine of excised patches from 2:1:1 POPC:POPE:POPG giant liposomes (black = WT, red = ELIC5, −60 mV holding voltage). The ELIC5 currents show sustained response to 10 mM cysteamine, followed by slow deactivation with the removal of cysteamine. **(b)** Inset that shows ELIC5 (red) single channel currents taken from the deactivation trace (i.e. channel closure) or WT (black) single channel currents in the presence of 10 mM cysteamine. **(c)** Electron density map of ELIC5 in 2:1:1 POPC:POPE:POPG nanodiscs + 10 mM cysteamine (grey) with outer leaflet lipid density (yellow) and black lines approximating the bilayer. **(d)** View of the pore lining M2 helix from the extracellular side showing 16’ (F247) and 9’ (L240) side chains from WT apo, WT CA (WT + 10 mM cysteamine), ELIC3 CA (ELIC3 + 10 mM cysteamine), and ELIC5 CA (ELIC5 + 10 mM cysteamine). All are structures in 2:1:1 POPC:POPE:POPG nanodiscs. **(e)** Ion permeation pathway as determined by HOLE^69^ of the same structures shown in (D). Labeled are 16’, 9’ and 2’ side chains which form the narrowest portions of the pore in different structures. **(f)** Pore radius as a function of distance along the pore axis for the same structures as (D) and (E).

**Table 1.**
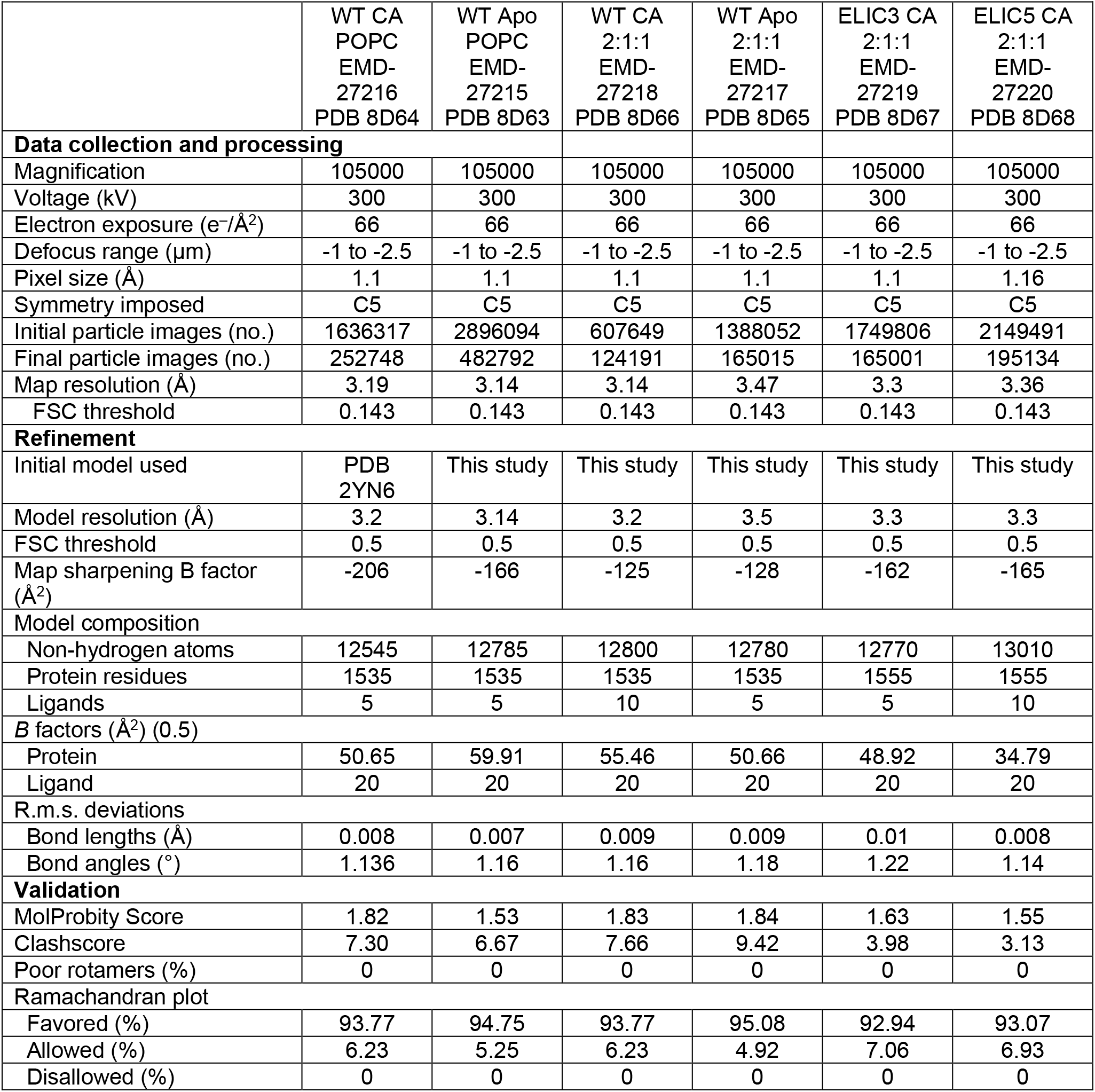
Cryo-EM data collection and refinement statistics

|  | WT CA<br>POPC<br>EMD-27216<br>PDB 8D64 | WT Apo<br>POPC<br>EMD-27215<br>PDB 8D63 | WT CA<br>2:1:1<br>EMD-27218<br>PDB 8D66 | WT Apo<br>2:1:1<br>EMD-27217<br>PDB 8D65 | ELIC3 CA<br>2:1:1<br>EMD-27219<br>PDB 8D67 | ELIC5 CA<br>2:1:1<br>EMD-27220<br>PDB 8D68 |
| --- | --- | --- | --- | --- | --- | --- |
| **Data collection and processing** |  |  |  |  |  |  |
| Magnification | 105000 | 105000 | 105000 | 105000 | 105000 | 105000 |
| Voltage (kV) | 300 | 300 | 300 | 300 | 300 | 300 |
| Electron exposure (e<sup>-</sup>/Å<sup>2</sup>) | 66 | 66 | 66 | 66 | 66 | 66 |
| Defocus range (μm) | -1 to -2.5 | -1 to -2.5 | -1 to -2.5 | -1 to -2.5 | -1 to -2.5 | -1 to -2.5 |
| Pixel size (Å) | 1.1 | 1.1 | 1.1 | 1.1 | 1.1 | 1.16 |
| Symmetry imposed | C5 | C5 | C5 | C5 | C5 | C5 |
| Initial particle images (no.) | 1636317 | 2896094 | 607649 | 1388052 | 1749806 | 2149491 |
| Final particle images (no.) | 252748 | 482792 | 124191 | 165015 | 165001 | 195134 |
| Map resolution (Å) | 3.19 | 3.14 | 3.14 | 3.47 | 3.3 | 3.36 |
| FSC threshold | 0.143 | 0.143 | 0.143 | 0.143 | 0.143 | 0.143 |
| **Refinement** |  |  |  |  |  |  |
| Initial model used | PDB<br>2YN6 | This study | This study | This study | This study | This study |
| Model resolution (Å) | 3.2 | 3.14 | 3.2 | 3.5 | 3.3 | 3.3 |
| FSC threshold | 0.5 | 0.5 | 0.5 | 0.5 | 0.5 | 0.5 |
| Map sharpening B factor (Å<sup>2</sup>) | -206 | -166 | -125 | -128 | -162 | -165 |
| **Model composition** |  |  |  |  |  |  |
| Non-hydrogen atoms | 12545 | 12785 | 12800 | 12780 | 12770 | 13010 |
| Protein residues | 1535 | 1535 | 1535 | 1535 | 1555 | 1555 |
| Ligands | 5 | 5 | 10 | 5 | 5 | 10 |
| **B factors (Å<sup>2</sup>) (0.5)** |  |  |  |  |  |  |
| Protein | 50.65 | 59.91 | 55.46 | 50.66 | 48.92 | 34.79 |
| Ligand | 20 | 20 | 20 | 20 | 20 | 20 |
| **R.m.s. deviations** |  |  |  |  |  |  |
| Bond lengths (Å) | 0.008 | 0.007 | 0.009 | 0.009 | 0.01 | 0.008 |
| Bond angles (°) | 1.136 | 1.16 | 1.16 | 1.18 | 1.22 | 1.14 |
| **Validation** |  |  |  |  |  |  |
| MolProbity Score | 1.82 | 1.53 | 1.83 | 1.84 | 1.63 | 1.55 |
| Clashscore | 7.30 | 6.67 | 7.66 | 9.42 | 3.98 | 3.13 |
| Poor rotamers (%) | 0 | 0 | 0 | 0 | 0 | 0 |
| **Ramachandran plot** |  |  |  |  |  |  |
| Favored (%) | 93.77 | 94.75 | 93.77 | 95.08 | 92.94 | 93.07 |
| Allowed (%) | 6.23 | 5.25 | 6.23 | 4.92 | 7.06 | 6.93 |
| Disallowed (%) | 0 | 0 | 0 | 0 | 0 | 0 |

Since POPG stabilizes the open state of ELIC, we aimed to capture an open conformation of ELIC using gain-of-function mutations. We began with the previously reported gain-of-function triple mutant V261Y/G319F/I320F (ELIC3)^29^ that introduces three aromatic residues at the top of the M4 helix (M4) (Extended Data Fig. 4a). As previously reported, ELIC3 caused a left-shift in the agonist dose response curve, slowed the rate of current decay in the presence of agonist, and slowed deactivation as assessed by excised patch-clamp recordings from 2:1:1 giant liposomes (Extended Data Fig. 4b-4e). Interestingly, the structure of agonist-bound ELIC3 in 2:1:1 nanodiscs, determined at 3.3 Å resolution (Extended Data Fig. 2, Table 1), showed opening of the pore at 16’ but was still occluded at 9’ (Fig. 2d-2f). Seeking a mutant that remains open in the presence of agonist, we added the mutations P254G (previously shown to slow deactivation^30^) and C300S (previously shown to slow current decay in the presence of agonist^12, 31^), and produced P254G/C300S/V261Y/G319F/I320F (ELIC5) (Extended Data Fig. 4a). Strikingly, ELIC5 showed a deactivation rate ∼1200x slower than WT and no evidence of current decay in the presence of 10 mM cysteamine (Fig. 2a, Extended Data Fig. 4e), indicative of profound stabilization of the open state. In four recordings of ELIC5 currents in response to 10 mM cysteamine where single channel openings were resolved, there was no evidence of channel closure for several minutes (Extended Data Fig. 4g), suggesting that ELIC5 has a high open probability at steady state. Consistent with its gain-of-function phenotype, a structure of ELIC5 in 2:1:1 nanodiscs with 10 mM cysteamine, determined at 3.4 Å resolution (Extended Data Fig. 2 and 5, Table 1), yielded an open pore (Fig. 2d-2f) with a pronounced outer leaflet phospholipid-shaped density (Fig. 2c).

The structures of WT, ELIC3 and ELIC5 reveal different pore conformations that indicate a probable sequence of changes from resting to open (Fig 2d and 2e). The WT apo structure shows constrictions at 16’ and 9’ (Fig. 2e), consistent with resting pore conformations reported for the nAchR^32–34^. In the presence of agonist, the top of M2 in WT shows a modest blooming of 16’ consistent with a pre-active conformation^28^. The ELIC3 gain-of-function mutant further opens at 16’ with the F247 side chain turning away from the pore into an intersubunit space, but otherwise resembles the WT pre-active conformation. Finally, in ELIC5, 9’ rotates away from the pore into an intersubunit space and 16’ opens further, such that the narrowest diameter of the pore is now at 2’ (Q233) (Fig. 2f). The pore diameter at 2’ measures 7.3 Å, which is consistent with the capacity of ELIC to conduct large quaternary amines^35^. ELIC5 has the same single channel conductance as WT (Fig. 2b, Extended Data Fig. 4f); this along with its high open probability strongly suggests that the ELIC5 structure represents a functionally-relevant open conformation. Hereafter, to examine POPG modulation of ELIC, we will focus on WT apo, WT agonist-bound (WT CA) and the ELIC5 agonist-bound (ELIC5 CA) structures all in 2:1:1 nanodiscs, which likely represent resting, pre-active and open conformations, respectively.

### ELIC opening is associated with an outer leaflet bound phospholipid

The ELIC5 CA structure shows a strong phospholipid density at an outer leaflet site within a groove formed by M4, M3 and M1 (Fig. 3a). The density has well-resolved features including nearly full density for the palmitoyl-oleoyl acyl chains (Fig. 3a). Of the six structures obtained in this study, three others show evidence of phospholipid density at this outer leaflet site, including WT apo structures in POPC and 2:1:1, and the WT agonist-bound structure in 2:1:1 (Extended Data Fig. 6a)^14^. Thus, among agonist-bound structures, phospholipid density at this site was not observed in POPC nanodiscs. To compare the strength of the phospholipid densities in 2:1:1 structures, we low pass filtered the maps to 3.5 Å resolution and displayed the map for each lipid at a sigma value of 3.0 (Fig. 3b). The maps show relatively weak lipid density in the WT CA structure compared to the ELIC5 CA structure, suggesting that a phospholipid stabilizes the open state relative to the pre-active state by binding to this site.

**Fig. 3.**
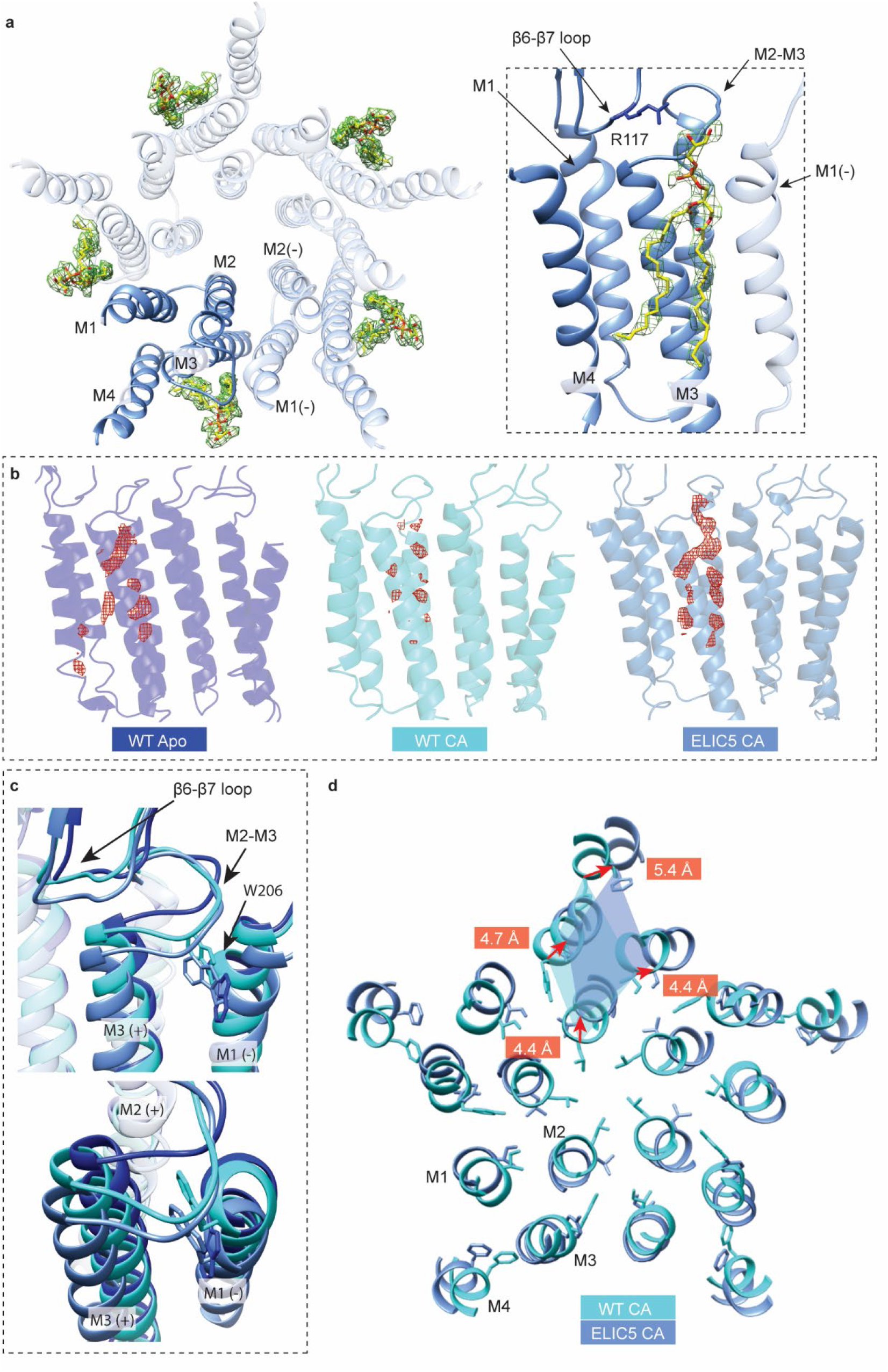
Outer leaflet phospholipid binding site and associated conformational changes. **(a)** *Left:* Structure of ELIC5 TMD viewed from extracellular side showing bound POPG (yellow) in the density map (green). *Right*: Side view of ELIC5 phospholipid binding site showing POPG (yellow) in the density map (green). The R117 side chain is shown. **(b)** Phospholipid density (red) displayed at σ-level of 3.0 from electron density maps of WT apo (resting), WT CA (pre-active), and ELIC5 CA (open). **(c)** *Top:* Conformational changes in the β6-β7 loop and the M2-M3 linker in WT apo, WT CA and ELIC5 CA. Shown is the W206 side chain for each structure. *Bottom*: Top down view of the conformational changes of the M2-M3 linker and W206 of WT apo, WT CA and ELIC5 CA. **(d)** View of the TMD helices from the extracellular side at the level of 9’ comparing WT CA and ELIC5 CA. Distances between C-α carbons of each structure are shown in red for each transmembrane helix.

The phospholipid binds at an outer leaflet site with the headgroup located below the β6-β7 loop and adjacent to the M2-M3 linker, and the diacylglycerol in a groove formed by M4, M3 and M1 (Fig. 3a). Of note, the mutated residues in ELIC5 are not located in this site (Extended Data Fig. 4a, Fig. 5a). Closer examination of this site shows major conformational changes between the ELIC5 CA, WT CA and WT apo structures. The β6-β7 loop progressively translates outward from WT apo to WT CA to ELIC5 CA (Fig. 3c), placing the basic residue, R117, less than 4 Å from the phospholipid headgroup in the ELIC5 CA open structure (Fig. 3a, Extended Data Fig. 6b). Movement of the β6-β7 loop is associated with an outward translation of the M2-M3 linker and the top of M3 and M1, which is largest in the ELIC5 CA structure (Fig. 3c). The outward translation of these structures is linked to a global outward blooming of 4-5 Å of all four transmembrane helices leading to opening of the pore (Fig. 3d). Thus, conformational changes in the phospholipid binding site of the open structure of ELIC, especially in the β6-β7 loop where R117 interacts with the phospholipid headgroup, reveals a plausible mechanism for stabilization of the open state by an anionic phospholipid such as POPG.

Activation of ELIC is also associated with movement of a semi-conserved aromatic residue, W206 in M1, which is implicated in binding of zwitterionic phospholipids^36^ and located in this outer leaflet phospholipid binding site (Fig. 3c). Indeed, we observed phospholipid density directly interacting with W206 in the WT apo structures in POPC and 2:1:1 nanodiscs (Fig. 3b, Extended Data Fig. 6b). However, while W206 faces into the bilayer in the resting structure, in agonist-bound structures (WT CA and ELIC5 CA), it flips into an intersubunit space displacing M3 and the M2-M3 linker and dramatically changing the structure of this lipid binding site (Fig. 3c, Extended Data Fig. 6b). We discovered that the tryptophan fluorescence of ELIC decreases with the addition of agonist, and that this decrease is entirely attributed to W206 (no change in fluorescence is observed in W206F with and without agonist, Extended Data Fig. 7a and 7b). Taking advantage of this unique agonist-dependent signal, we monitored tryptophan fluorescence of ELIC after rapid mixing of propylamine, to compare the rate of W206 movement with channel opening from the thallium flux assay. Although this measurement was performed with ELIC in DDM, the rate of decrease in tryptophan fluorescence was approximately 2-4x faster than channel opening in 2:1:1 or POPC liposomes from the thallium flux assay (Extended Data Fig. 7c). This finding indicates that W206 moves away from the protein-lipid interface during pre-activation, consistent with the WT CA structure being a pre-active conformation. W206 has been reported to form cation-pi interactions with phospholipids in the resting conformation favoring zwitterionic over anionic phospholipids^37^; however, movement of W206 away from the membrane with channel activation may alter the specificity of this site for phospholipids such as PG.

The WT apo, WT CA and ELIC5 CA structures show other notable differences in the ECD and TMD. In the ECD, agonist binding initiates movement of loop C towards the pore vestibule clamping down on the agonist (Extended Data Fig. 8a), a conformational change previously described in ELIC and other pLGICs^28^. However, in the ELIC5 CA structure, the agonist binding site remains clamped down but translates away from the pore (Extended Data Fig. 8a). Associated with this translation, there is a general expansion of the ECD β-sheets β1-β6 (Extended Data Fig. 3a). In the TMD, the outward blooming of the TM helices in the ELIC5 CA structure extends to the bottom of the transmembrane helices (Extended Data Fig. 8b), and is associated with outward translation and straightening of M4 below P305, a conserved proline that produces a kink in the M4 helix in non-conducting structures of ELIC (Extended Data Fig. 8c)^22^.

In summary, the ELIC5 CA structure shows key conformational changes associated with channel opening, and a putative mechanism by which a phospholipid stabilizes the open state of ELIC through an outer leaflet binding site.

### POPG modulates ELIC from the outer leaflet

To further assess the possibility that POPG modulates ELIC from this binding site in the outer leaflet, we measured ELIC function in asymmetric liposomes using the fluorescence stopped-flow flux assay^13, 25, 38–41^. ELIC was reconstituted in 3:1 POPC:POPE liposomes, after which the sample was split and treated with methyl-β-cyclodextrin alone (no-exchange), or methyl-β-cyclodextrin with POPG (exchange) set to introduce 25 mole% POPG to the outer leaflet (Fig. 4a)^39^. Outer leaflet POPG was assessed by zeta potential measurements^39^. The asymmetric proteoliposomes had a similar zeta potential as symmetric 2:1:1 POPC:POPE:POPG liposomes, indicating successful introduction of 25 mole% POPG in the outer leaflet and no significant flipping of POPG to the inner leaflet (Fig. 4b)^39, 42^. Moreover, the zeta potentials of asymmetric liposomes with 25 mole% outer leaflet POPG with and without ELIC were identical, indicating that ELIC does not cause flipping of POPG from the outer to inner leaflet or, itself, alter the zeta potential (Fig. 4b).

**Fig. 4.**
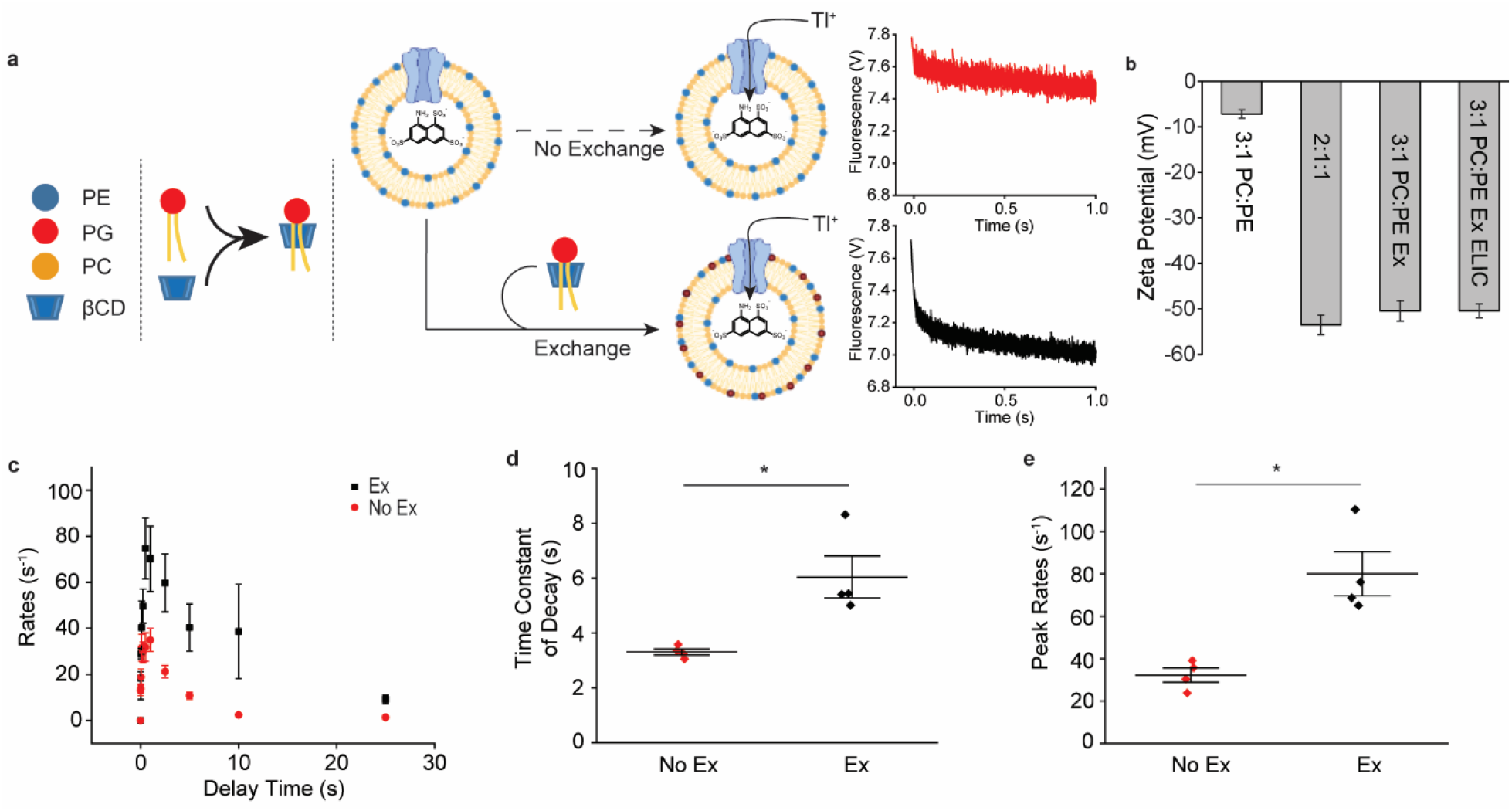
Asymmetric liposome stopped-flow Tl^+^ flux assay. **(a)** Schematic representation of asymmetric liposome stopped-flow Tl^+^ flux assay. Starting with 3:1 POPC:POPE liposomes reconstituted with ELIC, the sample was either kept as a symmetric proteoliposome and subjected to the Tl^+^ flux assay (top, “No Exchange”), or treated with mβCD/POPG to introduce 25 mole% POPG to the outer leaflet (bottom, “Exchange”). **(b)** Zeta potential measurements of symmetric 3:1 POPC:POPE liposomes (3:1 PC:PE), symmetric 2:1:1 POPC:POPE:POPG liposomes (2:1:1), asymmetric 3:1 POPC:POPE liposomes treated to introduce 25 mole% POPG to the outer leaflet (3:1 PC:PE Ex), and asymmetric 3:1 POPC:POPE proteoliposomes with ELIC treated to introduce 25 mole% POPG to the outer leaflet (3:1 PC:PE Ex ELIC) (n = 3, ± SEM). **(c)** Tl^+^ flux rates of WT ELIC in 3:1 PC:PE no exchange liposomes (red, “No Ex”) and exchanged asymmetric liposomes where 25 mol% POPG is introduced to the outer leaflet (black, “Ex”) (n = 4, ± SEM). Channel activation was elicited by 10 mM propylamine. **(d)** Weighted time constant of decay in channel activity in the presence of 10 mM propylamine from Tl^+^ flux assay comparing no exchange and exchange conditions (n = 4, ± SEM). * indicates p<0.05. **(e)** Peak rate of Tl+ flux in response to 10 mM propylamine in no exchange and exchange conditions (n = 4, ± SEM). * indicates p<0.05.

The addition of methyl-β-cyclodextrin alone did not alter ELIC responses. However, introduction of 25 mole% POPG to the outer leaflet increased the peak agonist response approximately two-fold and slowed the rate of decay in activity two-fold (Fig. 4c-4e). This effect qualitatively recapitulates the effect of POPG in symmetric liposomes (Fig. 1). Of note, the reconstitution efficiency and orientation of ELIC in the no-exchange and exchange liposomes are the same, since both samples are taken from the same batch of proteoliposomes. Thus, in the asymmetric liposome experiments, peak responses between no-exchange and exchange samples can be attributed to channel activity (e.g. open probability). Overall, these data definitively show that POPG modulates ELIC from the outer leaflet, increasing peak channel activity (i.e. gating efficacy) and slowing the decay in channel activity. Not having examined the effect of POPG in the inner leaflet alone, we cannot exclude the possibility of inner leaflet POPG modulation^13^.

### POPG binds specifically to the outer leaflet site in the open structure

Having established that POPG increases ELIC gating efficacy from the outer leaflet, we further assessed whether this effect is mediated by the phospholipid bound to the outer leaflet site. We hypothesized that this site is specific for POPG and that POPG binding favors the open over the pre-active conformation. To test this, we carried out streamlined alchemical free energy perturbation (SAFEP^24^) simulations to determine the relative affinities of POPC, POPE, and POPG for binding modes corresponding to the phospholipid-shaped density in WT CA (pre-active) and ELIC5 CA (open) structures. Representative density of POPG bound to the ELIC5 CA structure is shown in Figure 5A. Using SAFEP, we determined the relative probabilities of POPG, POPC, or POPE for the outer leaflet site, conditional upon the requirement that the glycerol backbone for each phospholipid remains within 6 Å of the reference configuration, which is the upper limit observed in simulations of the bound state (Supplementary Figure 1). These calculations revealed preferential binding of POPG for the outer leaflet site in a 2:1:1 POPC:POPE:POPG membrane for both WT CA and ELIC5 CA structures.

**Fig. 5.**
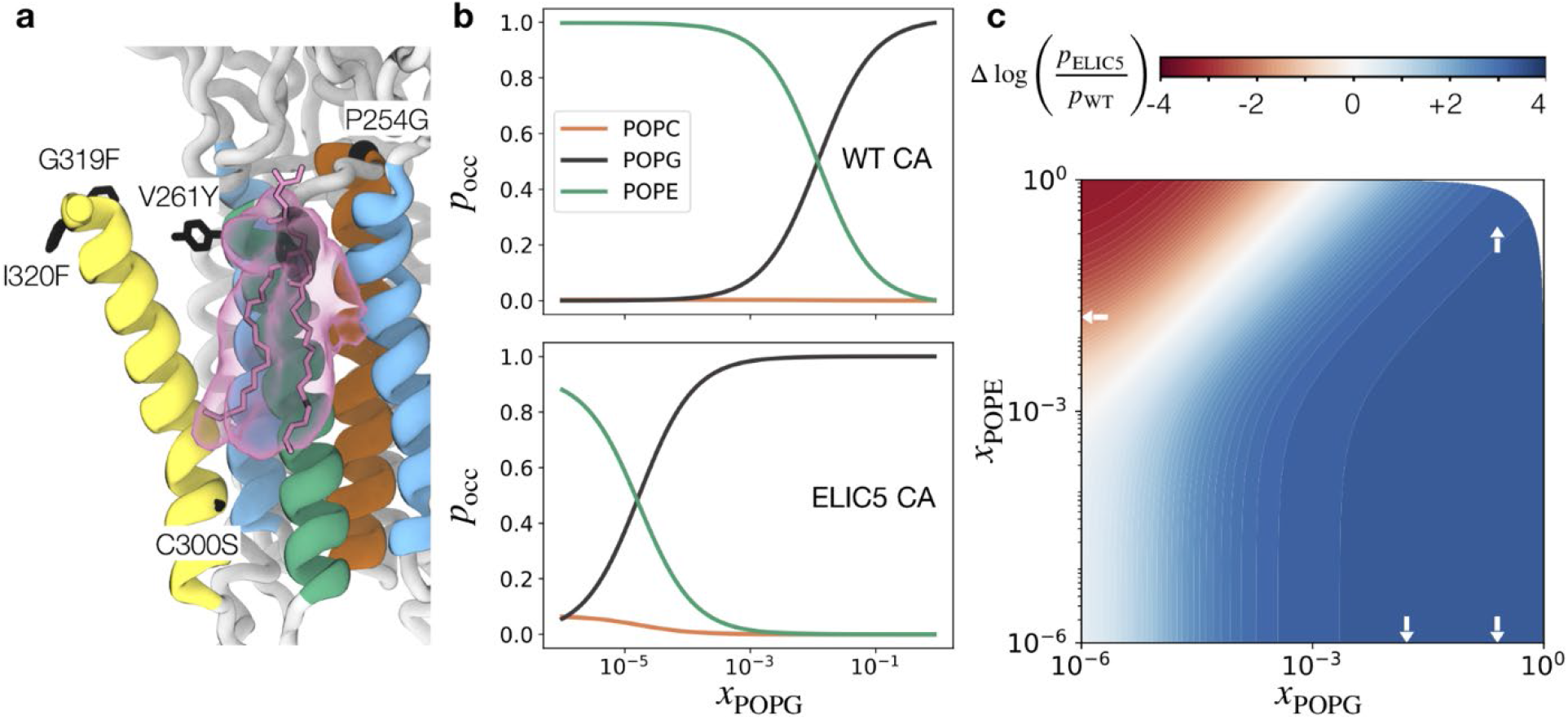
Predicted lipid occupancy for the outer leaflet phospholipid binding site. **(a)** ELIC5 CA open structure showing lipid density from 20 ns of equilibrium MD simulation compared to the reference structure modeled with POPG (pink licorice). The region of glycerol backbone density is shown in black, while the remaining lipid density is shown in pink. The five mutations of ELIC5 are shown in black licorice. (**b)** The SAFEP-calculated probability (p_occ_) that the structurally-identified site will be occupied by each of three possible lipids in a 2_POPC_:1_POPE_:*x*_POPG_ mixture, for both the WT CA conformation (top) and the ELIC5 CA conformation (bottom). **(c)** The predicted relative conformational stability 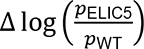 as a function of mole fraction of POPG and POPE in a POPG:POPE:POPC mixture. Red and blue correspond to greater stability of the WT CA and ELIC5 CA conformations, respectively. White arrows indicate compositions where bulk calculations were carried out; remaining values were extrapolated as described in Methods. Data in panels b and c represent over 3 μs of simulation; pharmacological models relied on SAFEP-calculated parameters as described in Methods.

To investigate state dependence and the effect of membrane composition, we extrapolated to a range of ternary compositions to obtain phospholipid binding curves (Fig. 5B). In a 2_POPC_:1_POPE_:*x*_POPG_ ternary mixture, POPC rarely occupies the outer leaflet site, while POPE occupies the site at very low *x*_POPG_. At higher *x*_POPG_, POPG displaces POPE from the site, indicating preferential binding of POPG for the outer leaflet site. We define the *x*_50_ as the mole fraction of POPG required to achieve 50% occupancy. From the curves in Figure 5B, we estimate that *x*_50_ ∼ 10^-5^ and *x*_50_ ∼ 10^-2^ for the ELIC5 and WT conformations, respectively, which is consistent with POPG stabilization of the ELIC5 CA conformation in these ternary mixtures. Estimated relative stabilities of ELIC5 CA and WT CA are shown for a continuum of ternary compositions in Figure 5C. Relative to pure POPC, where the relative stabilization is 0 by definition, adding POPE stabilizes the non-conducting WT CA conformation, and adding POPG stabilizes the open ELIC5 CA conformation.

Since the structure of the lipid binding site dramatically differs between apo and agonist-bound structures especially due to movement of W206 (Extended Data Fig. 6b), we examined POPG binding to this site in the WT apo structure. Equilibrium MD simulations of POPG in the binding mode observed in the WT apo structure showed unbinding of the POPG headgroup after about 10 ns of simulation time, consistent with previous simulations in apo ELIC structures (Extended Data Fig. 9)^37^. Thus, the outer leaflet site appears to only accommodate POPG in agonist-bound states. The SAFEP results predict that POPG occupation of this site stabilizes the open state of the channel.

### Functional role of the outer leaflet POPG binding site

We next tested the effect of mutations that directly interact with the phospholipid in the open structure (R117, T259), or are near this site and previously shown to interact with cardiolipin (S202, R318A, T259A and W206F) (Fig. 6a)^14^. The mutations were tested as for WT by examining the effect of 25 mole% outer leaflet POPG in paired (no exchange versus exchange) samples (Extended Data Fig. 10).

**Fig. 6.**
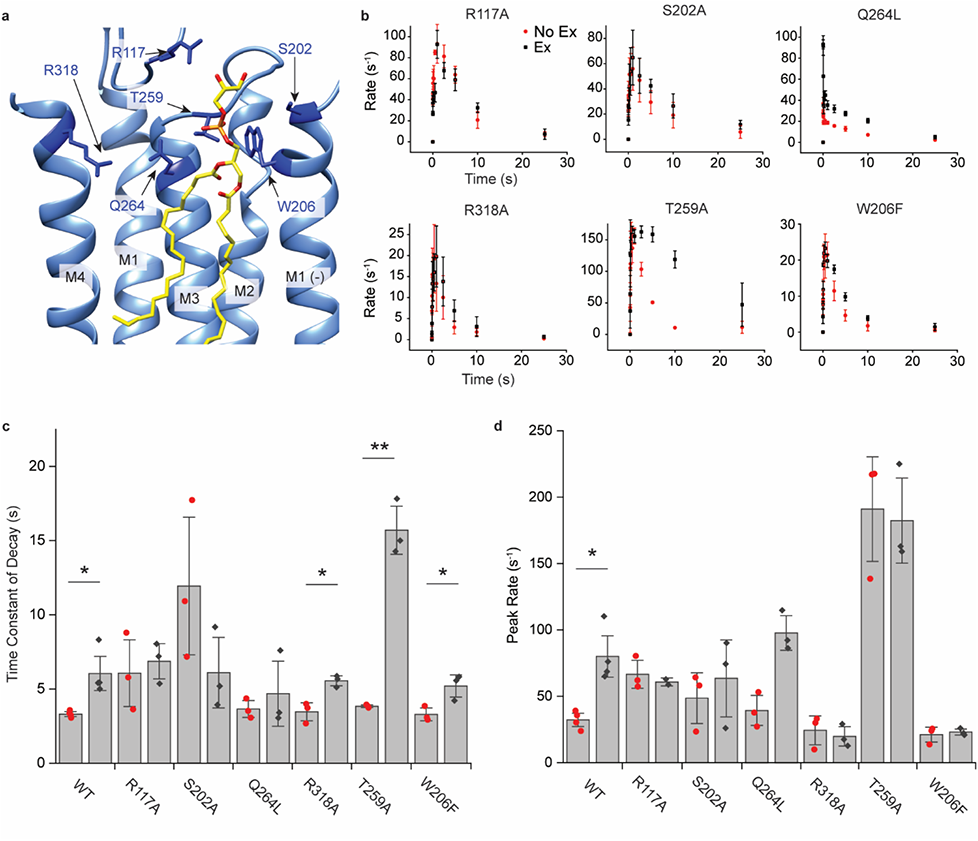
Mutational analysis of the outer leaflet phospholipid binding site. **(a)** ELIC5 CA open structure showing POPG (yellow) at the outer leaflet site and residues (dark blue) targeted by mutagenesis. **(b)** Tl^+^ flux rates of WT and mutants in response to 10 mM propylamine for no-exchange (red) and exchange (black, 25 mole% POPG in the outer leaflet) conditions (n = 3, ± SEM). **(c)** Weight time constant of decay in channel activity in response to 10 mM propylamine for no-exchange (red) and exchange (black) conditions (n = 4 for WT, n=3 for all mutants, ±SEM). **(d)** Same as (c) showing peak rates of Tl^+^ flux in response to 10 mM propylamine for no-exchange (red) and exchange (black) conditions (n = 4 for WT, n=3 for all mutants, ±SEM). * indicates p<0.05. ** indicates p<0.01.

Three mutations (Q264L, R117A, and T259A) showed an apparent loss-of-function in that agonist responses were low (T259A) or undetectable (Q264L and R117A); to improve the signal, we increased the amount of protein 10x for the reconstitution of these mutants. Despite the mutations having various effects on ELIC agonist responses, comparison of no exchange and exchange samples allowed direct assessment of the effect of POPG. The mutations can be categorized into those where both effects of POPG (increased peak response and slowed decay rate) are abolished (R117A, S202A), those where POPG modulation is preserved (Q264L), and those where POPG only slows the rate of decay in channel activity (R318A, T259A, W206F) (Fig. 6b and 6c). R117A, which mutates a basic residue near the phospholipid headgroup in the open structure (Fig. 6a), strikingly ablates both measures of POPG modulation (Fig. 6c and 6d). S202A at the top of M1 also produced a loss of POPG modulation. Q264L activation and decay in channel activity was very fast such that maximum activity was seen at the shortest time point (i.e. 10 ms delay time), making the exact peak response and rate of decay uncertain. Nevertheless, Q264L showed a 2x increase in the measurable peak response with POPG (Fig. 6c) and significantly higher activity at delay times of 2.5 to 10 s with POPG (Fig. 6b), indicating that POPG positively modulates peak response and decay in activity in this mutant. Interestingly, Q264 interacts with the phospholipid headgroup (distance of 2.5 Å) in the resting structure (Extended Data Fig. 6b), but rotates away to face M4 in both agonist-bound pre-active and open structures > 7 Å removed from the phospholipid headgroup (Fig. 6a and Extended Data Fig. 6b). Thus, the preserved modulatory effect of POPG in Q264L is consistent with the notion that POPG modulates the equilibrium between agonist-bound states; in the pre-active and open conformations, POPG at the proposed binding site does not closely interact with Q264.

R318A, T259A and W206F abolished the effect of POPG on peak response, but still showed an increase in the time constant of decay in channel activity with POPG, with some variability (Fig. 6b-6c). The reason for this discrepancy, especially T259A which showed a profound slowing of decay in channel activity with POPG, is unclear (Fig. 6c). It may reflect the fact that POPG effects on peak response and decay in activity result from changes in the relative stability of distinct non-conducting states such as pre-active and desensitized states, respectively. Without a structure in a desensitized conformation, it is difficult to predict the impact of these mutations on phospholipid binding. Overall, the mutations have complex effects on channel function and POPG modulation. However, the unequivocal effect of R117A on POPG modulation, which disrupts a charged interaction with the POPG headgroup, and the loss of POPG effect on peak response in multiple mutations implicates the outer leaflet M4-M3-M1 site in the mechanism of POPG modulation.

## DISCUSSION

It has been known for four decades that anionic phospholipids are critical for maximal agonist response in the nAChR^9, 43^. More recently, anionic phospholipids have been shown to modulate ELIC^13, 14^. Many structures of different pLGICs including ELIC show bound phospholipids^19–21^, intimating that phospholipid modulation is mediated by direct binding to a specific site(s). The results of this study establish a structural mechanism for leaflet-specific, site-specific POPG modulation of the pLGIC, ELIC. POPG binds favorably to an outer leaflet site between M4, M3, and M1, in the open conformation of the channel. A recent structure of ELIC in SMA nanodiscs showed cardiolipin bound to a similar site^14^. Like POPG, cardiolipin also positively modulates ELIC. Another study using MD simulations argued that zwitterionic lipids such as PC selectively bind to this site forming a cation-pi interaction with W206^36^. However, both studies examined resting state structures where W206 faces into the bilayer. In our agonist-bound structures including the open structure, W206 has turned into an intersubunit space, drastically changing the structure of this lipid binding site and making PG occupancy much more favorable. Therefore, it seems plausible that cardiolipin stabilizes the open state of ELIC also by binding to this site, but with a different binding mode than what was observed in the resting structure^14^. Interestingly, a recent study demonstrated that the polyunsaturated fatty acid, docosahexaenoic acid (DHA), inhibits ELIC by binding to a site overlapping with the POPG site. DHA produces the opposite effect as POPG: it decreases peak agonist response and increases the rate of decay in activity^12^. Thus, it appears that competition by different lipids (i.e. anionic phospholipids and fatty acids) for this site modulates the relative stability of open and agonist bound non-conducting states in ELIC.

Using asymmetric liposomes, we definitively establish that POPG modulates ELIC from the outer leaflet. This finding is agreeable with the fact that PG is enriched in the outer leaflet of gram negative bacteria membranes where ELIC is expressed^44^. Whether anionic lipid species also modulate eukaryotic pLGICs through this outer leaflet binding site remains to be determined. However, lipid densities have been observed in the structures of the muscle nAchR, GluCl and the 5-HT_3A_R at this outer leaflet site^19, 32, 33, 45^. In addition, MD simulations of the glycine receptor showed state-dependent binding of phospholipids between inactive and active conformations at the top of M3 and M1—a site analogous to the POPG binding site in ELIC^46^. Interestingly, R117 in the β6-β7 loop, which forms an electrostatic interaction with the PG headgroup in the open structure, is conserved in GLIC and human pLGICs (Supplementary Fig. 2). Thus, there is structural and computational evidence to suggest that a mechanism, similar to that proposed in the present study, exists in human pLGICs. Anionic phospholipids are localized to the inner leaflet of mammalian plasma membranes^47^; yet, it is thought that some anionic phospholipids such as phosphatidylserine exist in the outer leaflet of neuronal membranes and are dynamically regulated^48^. It is possible that anionic phospholipids in the outer leaflet fine-tune agonist responses of mammalian pLGICs through this site, or that other lipids more abundant in the outer leaflet act through this site.

We report the first open structure of ELIC; all prior structures of ELIC show a resting conformation, with the exception of an agonist-bound structure in POPC nanodiscs displaying partial activation^14, 22, 28, 49, 50^. While we cannot exclude the possibility that the ELIC5 structure is an artifact of mutations, the fact that this mutant has a high steady-state open probability in the presence of agonist shows excellent agreement between structure and function. The conformational changes transitioning from resting to pre-active to open in ELIC are generally conserved in other pLGICs, although our structures reveal marked differences between pre-active and open states that are not appreciated to the same extent in other pLGICs structures^19, 33, 34, 51, 52^. These include a significant outward blooming of the TM helices and expansion of the ECD β-sheets (β1-β6), indicating that the transition from pre-active to open involves global conformational changes spanning the ECD and TMD. Such an observation is consistent with a model that long-range global allosteric rearrangements underlie the transitions between functional states in pLGICs.

In conclusion, we present the first open structure of ELIC and a structural mechanism for positive allosteric modulation of ELIC by the anionic phospholipid, POPG. POPG specifically binds to an outer leaflet M4-M3-M1 site, stabilizing outward movement of the β6-β7 loop and M2-M3 linker, and opening of the pore.

## METHODS

### Protein mutagenesis, expression, and purification

The pET-26-MBP-ELIC was provided by Raimund Dutzler (Addgene plasmid # 39239) and was used for the generation of both WT and mutant ELIC. Site-directed mutagenesis was conducted using the Quickchange method and subsequently confirmed by Sanger sequencing (Genewiz, Plainfiled, NJ). WT and mutant ELIC were expressed as previously reported^15, 49^ in OverExpress C43 (DE3) *E. Coli* (Lucigen, Middleton, WI). Terrific Broth (Sigma, St. Louis, MO) was used to grow cultures which were induced with 0.1 mM IPTG for ∼16 h at 18°C. The cells were pelleted, resuspended in Buffer A (20 mM Tris pH 7.5, 100 mM NaCl) with complete EDTA-free protease inhibitor (Roche, Indianapolis, IN), and lysed using an Avestin C5 emulsifier at ∼15,000 psi. Membranes were pelleted by ultracentrifugation, resuspended in Buffer A, solubilized with 1 % DDM (Anatrace, Maunee, OH), and incubated with amylose resin (New England Biolabs, Ipswich, MA) for 2 h. The amylose resin was washed with 20 bed volumes of Buffer A containing 0.02% DDM and 0.05 mM TCEP, and eluted with 40 mM maltose. The eluted protein was digested with HRV-3C protease (Thermo Fisher, Waltham, MA) (10 units per mg ELIC) overnight at 4°C then purified over a Sephadex 200 Increase 10/300 (GE Healthcare Life Sciences, Pittsburgh, PA) size exclusion column in Buffer A with 0.02% DDM.

### Nanodisc reconstitution

Phospholipids (POPC, POPE, POPG) (Avanti Polar Lipids, Alabaster AL) were used as received or combined to the desired ratio in chloroform, then dried under a stream of nitrogen and placed in a vacuum desiccator overnight. The lipids were then rehydrated in Buffer A to a concentration of 7.5 mg/ml (∼10 mM), sonicated for 5-10 minutes, and subjected to 5-10 freeze-thaw cycles using liquid nitrogen and a 37°C heat block, and finally extruded with a 400 nm filter 40x (Avanti Polar Lipids, Alabaster, AL). Samples were prepared using a 1:2:200 ratio of ELIC:MSP1E3D1:phospholipid. 300 µl samples were prepared by combining 74 µL Buffer A, 30 µL of ∼10 mM liposomes, and 70 µL of 1% DDM in Buffer A yielding a 0.23% DDM solution which was incubated at RT for 3 h. Next, 93 µL of 3 mg/mL (∼16 µM) ELIC, or ELIC mutant, was added with 33 µL of MSP1E3D1 at 3 mg/mL, then incubated at RT for 1.5 h. MSP1E3D1 was expressed and purified according to previous reports, without removal of the His-tag^53^. The pMSP1E3D1 plasmid was a gift from Stephen Sligar (Addgene plasmid # 20066). Following incubation, the samples were cooled on ice for 10 min followed by the addition of 80 mg Bio-beads SM-2 Resin (Bio-Rad, Hercules, CA), then rotated at 4°C overnight. The nanodisc solution was purified on a Sephadex 200 Increase 10/300 column using 10 mM HEPES with 100 mM NaCl at pH 7.5.

### Cryo-EM sample preparation and imaging

3.0 µl of purified ELIC nanodiscs at an approximately concentration of 1.2 mg/ml were pipetted onto Quantifoil R2/2 copper grids post plasma cleaning in an H_2_/O_2_ plasma for 60 s (Solaris 950. Gatan, Warrendale, PA). Agonist bound samples were prepared by combining the purified channels with cysteamine to a final agonist concentration of 10 mM. Post application of the sample, the grid was blotted for 2 s at ∼100% humidity and vitrified via plunge freezing into a solution of liquid ethane using a Vitrobot Mark IV (ThermoFisher Scientific, Prague, Czech Republic). Post vitrification, grids were mounted in Autogrid holders and transferred to a Titan Krios 300 kV Cryo-EM (ThermoFisher Scientific, Eindhoven, Netherlands). Datasets were collected in an automated fashion with the EPU acquisition software using a Gatan K2 Summit camera mounted on a GIF BioQuantum 968 energy filter (Gatan, Warrendale, PA). Movies were collected in counting mode with a nominal pixel size of 1.1 Å with the GIF set to a slit width of 20 eV operating in zero-loss mode. Dose fractionated movies were acquired over a defocus range of −1 to −2.5 µm. Each movie consisted of 40 individual frames with a per-frame exposure time of 200 ms, resulting in a dose of 66 electrons per Å^2^.

### Single particle analysis and model building

Raw movies were motion corrected and subjected to dose weighting with MotionCor2^54^. Motion corrected and dose weighted images were subject to contrast transfer function (CTF) determination using GCTF^55^. After CTF determination, low-quality micrographs were manually discarded. For each dataset ∼2500 particles were hand selected for 2D class averaging from which templates were derived for automated particle picking in RELION3^56^. Particles were extracted and subjected to multiple rounds of 2D and 3D classification. For WT ELIC CA in POPC nanodiscs, an initial 3D model was generated using a 40 Å-low-pass filtered map of PBD 2YN6 and used for subsequent 3D classifications. The other five datasets utilized a 40 Å-low-pass filtered map from the WT ELIC CA POPC dataset as an initial model. 3D classification was performed using C5 symmetry and no significant differences were identified among good classes. The best 3D refined maps were then masked and post-processed, followed by CTF refinement and Bayesian polishing. To compare the strength of the phospholipid density between WT apo, WT CA and ELIC5 CA structures, each structure was low-pass filtered to 3.5 Å^57^. Next, the lipid density was visualized in PyMol by adjusting the σ-level to 3.0^58^. Other images of the structures were generated using UCSF Chimera^59^.

An initial model of the WT CA 2:1:1 structure was generated by docking an ELIC crystal structure (PDB: 2YN6) into the cryo-EM density map using UCSF Chimera and then performing real space refinement using PHENIX^59, 60^. The remaining structure was manually built *de novo* into the density map using COOT prior to iterative real space refinement using PHENIX^61^. Prior to finalizing the model manual adjustments were made using COOT and verified in PHENIX. The other structures were built by docking the WT CA 2:1:1 structure into each density map and following the same procedure above. A non-proteinaceous density was observed in the agonist binding site of structures obtained in the presence of 10 mM cysteamine. Cysteamine was placed in this density in an orientation previously shown to be favorable by MD simulations^28^. Resolved phospholipid-like densities (a head group and two branching tails) were fit with POPG for those structures determined in 2:1:1 nanodiscs and POPC for those structures determined in POPC nanodiscs. Phospholipids were generated using ELBOW in PHENIX.

### Giant liposome excised patch-clamp recordings of ELIC

Giant liposome formation and excised patch-clamp recordings of ELIC were performed as previously described^12, 13^. 0.5 mg of WT ELIC, ELIC3 or ELIC5 were reconstituted in 5 mg of 2:1:1 POPC:POPE:POPG liposomes, using DDM destabilization and biobeads. 10 mM MOPS pH 7 and 150 mM NaCl (MOPS buffer) was used throughout. Proteoliposomes were pelleted and resuspended in ∼80 µl, and frozen in 10 µl aliquots. Giant liposomes were formed by dehydrating 10 µl of proteoliposomes on a glass coverslip in a vacuum desiccator for ∼20 min followed by rehydration with 100 µl of MOPS buffer overnight at 4 °C and 2 hr at RT the following day. The giant liposomes were resuspended by pipetting and applied to a recording chamber. The liposomes were allowed to settle for 15 min, after which free liposomes and debris were removed by exchanging the bath solution. Bath and pipette solutions contained MOPS buffer and 0.5 mM BaCl_2_. The addition of BaCl_2_ was necessary to facilitate the formation of giga-ohm seals. However, this concentration of BaCl_2_ also leads to a right-shift in the agonist dose-response curve, such that the EC_50_ for cysteamine is ∼4-5 mM instead of 1-2 mM. Of note, no BaCl_2_ was used in the fluorescence stopped-flow liposomal flux assay, such that the dose response curve for propylamine in 2:1:1 liposomes yielded an expected EC_50_ of ∼2 mM. All recordings were voltage-clamped at −60 mV and data collected at 5 kHz using an Axopatch 200B amplifier and Digidata 1440A (Molecular Devices, San Jose, CA). Rapid application or washout of agonist were performed using a three-barreled flowpipe mounted to and controlled by a SF-77B fast perfusion system (Warner Instrument Corporation, Hamden, CT). The 10-90% exchange time was previously determined to be ∼2 ms in this system using a liquid junction current across an open pipette, such that solution exchanges are sufficiently fast to capture ELIC activation and deactivation kinetics. Deactivation time courses were fit with a single exponential while the time course of current decay in the presence of agonist was fit with a double exponential. Weighted time constants were calculated using:

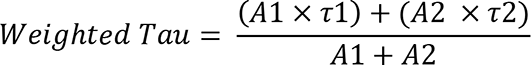

where A1 and A2 are the amplitudes of the first and second exponential components.

### Stopped-flow fluorescence recordings

The fluorescence sequential-mixing stopped-flow liposomal flux measurements were carried using an SX20 stopped-flow spectrofluorometer (Applied Photophysics, Leatherhead, UK) as previously described^13, 25^. 7.5 mg of the respective lipid mixture (POPC, 1:1 POPC: POPE, 1:1 POPC:POPG, 3:1 POPC:POPE, 3:1 POPC:POPG, 2:1:1 POPC:POPE:POPG) was dried in a glass vial to a thin film under a constant stream of N_2_. The lipids were then further dried under vacuum in a desiccator overnight. The following day the lipids were hydrated in 500 µl of reconstitution buffer (100 mM NaNO_3_, 10 mM HEPES, pH 7) (Buffer R), 250 µl of 75 mM 8-Aminonaphthalene-1,3,6-Trisulfonic Acid (ANTS) in buffer R, and doped with ∼18 mg of CHAPS. The mixture was vortexed and then sonicated with heat until completely solubilized. Following the sonication, the solution was brought to RT then 3.5 µl of 6.7 mg/ml ELIC in Buffer A with 0.02% DDM was added (3 µg per mg of lipid). This combined solution was incubated at RT for 30 min, followed by the addition of 1 ml of 1:1 v/v SM-2 BioBeads in Buffer R and an additional 250 µl of 75 mM ANTS in Buffer R. This mixture was incubated at RT for 2.5 hr with gentle mixing. The liposome suspension was then extruded with an Avanti mini-extruder and 0.1 µm membrane (30x). The extruded liposomes were stored at 13°C overnight with gentle rotation. The next day, the liposomes were diluted to 2.5 mL using assay buffer (140 mM NaNO_3_, 10 mM HEPES, pH 7) (Stopped-flow Buffer A), loaded into a PD-10 desalting column (Cytiva, Marlborough, MA), eluted with 3 mL of Stopped-flow Buffer A and subsequently diluted 5-fold with Stopped-flow Buffer A. Using propylamine as the agonist, the assay and analysis was carried out as previously described13,25. The time course of decay in channel activity was also fit to a double exponential and a weighted time constant was calculated as was done for the patch-clamp data.

### Tryptophan stopped-flow fluorescence measurements

To measure the tryptophan (Trp) fluorescence emission of ELIC, WT or W206F ELIC was diluted to 3 ml (0.02 mg/ml) and an emission spectrum was collected in a quartz fluorescence cell using a Fluoromax Plus Spectrofluorometer (Horiba Scientific, Piscataway, NJ). Excitation was set at 295 nm and an emission spectrum was collected from 320 to 380 nm. Trp fluorescence measurements with rapid mixing of agonist was carried out using the SX20 stopped-flow spectrofluorometer, outfitted with a 295 nm LED excitation source. Purified ELIC was diluted to 0.1 mg/ml in Trp Stopped-flow Buffer A (10 mM Tris pH 7.5, 100 mM NaCl, 0.05% DDM). A single mixing experiment was performed by mixing ELIC with equal volumes of Buffer A + propylamine. Propylamine was prepared at varying concentrations at 2x the final concentration. The instrument dead time was ∼1 ms, enabling capture of most of the time course of change in Trp fluorescence. The time course of decrease in Trp fluorescence after rapid mixing with propylamine was fit to a single exponential to obtain a time constant for the rate of decrease and a measure of the extent of decrease.

### Asymmetric liposome formation and analysis

Acceptor liposomes were prepared as described in the **Stopped-flow fluorescence recordings** section until completion of the extrusion step. The preparation was scaled up 2x to produce samples for the no-exchanged and POPG-exchanged conditions. After extrusion and storage of the liposomes overnight at 13°C, the double liposome preparation was split into two equal volume aliquots of ∼1.5 ml. Using the excel spreadsheet provided by Markones et al^39^, donor POPG and methyl-β-cyclodextrin (MβCD) concentrations and volumes were calculated to produce an outer leaflet POPG content of 25 mole% in the liposomes. The calculations assumed a ∼6.66 mM lipid concentration and 1.5 mL volume of the acceptor liposome solution^39^. A MβCD-POPG (50% saturation of MβCD by POPG) solution was formed by combining 224 µL 7.5 mM POPG in Stopped-flow Buffer A, 650 µl 300 mM MβCD in Stopped-flow Buffer A, and 126 μL Stopped-flow Buffer A, which was then mixed at 50°C and 1000 RPM for 20 min using an Eppendorf thermomixer. In parallel, an analogous POPG-free solution was generated and treated in the same manner except 224 µL of Stopped-flow Buffer A was used *in lieu* of the POPG solution. Critically, the solutions and thermomixer were cooled to 28°C. Following cooling, the 1.5 ml of acceptor liposomes were combined with the respective 1 ml MβCD solution, either containing POPG (exchange condition) or no POPG (no exchanged condition). These samples were mixed at 28°C and 400 RPM for 20 min using a thermomixer to complete the transfer of POPG to the liposomes. After mixing, the ∼2.5 ml of each respective sample were loaded on a PD-10 column and eluted with 3 mL Stopped-flow Buffer A. The fluorescence stopped-flow liposomal flux assay and analysis of the data were carried out on both the no exchange and exchange samples in sequence as previously described^13^.

### Zeta potential measurements

Zeta potential (ζ) measurements were made on a Malvern Zetasizer Nano-ZS ZEN 3600 using a flow-through high concentration zeta potential cell (HCC) following a previously described method^39, 40^. The instrument was equilibrated for 20 min at 28°C prior to analysis. Instrument settings included: temperature 28°, 90 s incubation time, 10-100 runs, and the dispersant settings were set as water for all samples. ζ was measured in triplicate for each sample. Prior to each measurement the HCC was flushed with copious volumes of preheated ddH_2_O water and dried under a flow of N_2_. The sample was prepared using liposomes immediately after desalting on the PD-10 column, and diluting 100 µl of liposomes with 100 mM HEPES at pH 7 to a final volume of 1 ml (final MβCD ∼ 4.3 mM). The sample was then loaded into a 1 ml syringe and the liposome suspension was driven into the cell until all air was displaced. Upon each subsequent measurement an additional ∼250 µl was pushed through the cell to displace old solution. After use, the HCC was disassembled and cleaned with 1% Hellmanex II in ddH_2_O and dried^39^.

### Molecular Dynamics Simulation

Initial protein-lipid complexes were modeled directly from cryo-EM maps with lipid densities. The first 11 protein residues were not modeled. The lipid densities were modeled using POPC, POPE, or POPG. These initial structures were then inserted into POPC membranes and equilibrated using default procedures from CHARMM-GUI^62^. All simulations were carried out using the CHARMM36m forcefield^63^ with cation-pi corrections^64^ Simulations were executed using NAMD 2.14^65^ in a semi-isotropic NPT ensemble tailored for membranes: a Langevin piston barostat with a target temperature of 303.15 K and a target pressure of 1 atmosphere with isotropic fluctuations enforced in the membrane plane. The piston period was increased to 200 steps (2 fs time step) and the decay to 100; this reduced instabilities due to higher frequency fluctuations. Following the default relaxation scheme of CHARMM-GUI^62, 66^, systems were further simulated for at least 20 more nanoseconds to ensure equilibration and to sample the unbiased NPT ensemble. Membrane-only systems were similarly created and equilibrated.

### Streamlined Alchemical Free Energy Perturbation (SAFEP) Simulations

We performed binding free energy estimation using the Molecular Dynamics Simulation settings previously described, with alchemical calculations using the SAFEP framework^24^. SAFEP is a simplified approach to calculating free energies of ligand binding using free energy perturbation (FEP). A primary challenge in FEP calculations is preventing spontaneous unbinding during simulation which would adversely impact the sampling efficiency. In the SAFEP framework^24^, the bound state is defined as a region of the configuration space for the ligand (RMSD) relative to the bound pose in the protein frame of reference (a.k.a. the “Distance-from-Bound-Configuration” or DBC). Interleaved-double wide sampling (IDWS) improves efficiency of any FEP by sampling in both the forward and backward lambda directions alternately (unlike traditional FEP in which each lambda value is simulated twice; once with forward sampling and once with backward sampling).

As in other relative FEP methods, the zero-sum nature of thermodynamic cycles is exploited to calculate interesting relative free energy differences from more computationally accessible data. In this case, the alchemical free energies of transformation calculated by SAFEP (summarized in Table 2) can be combined to calculate relative free energies of replacement of one lipid with another:

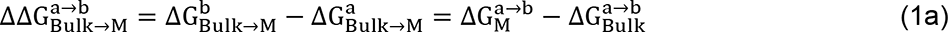

**Table 2.**
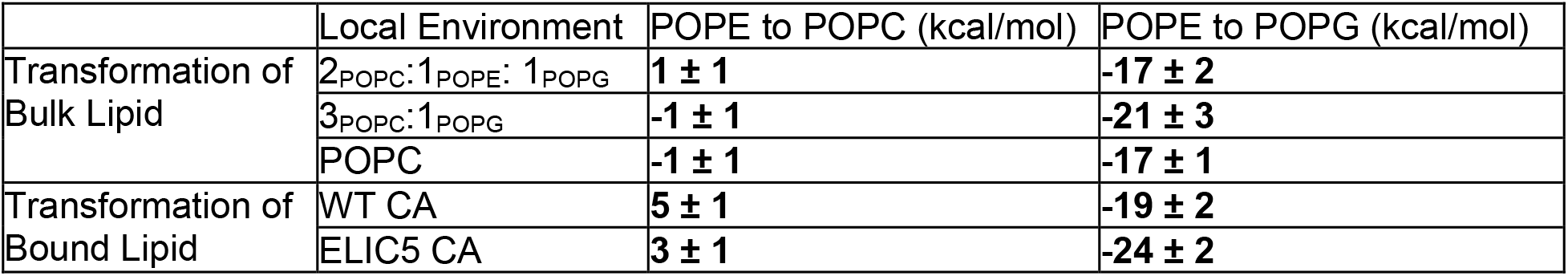
Free energy differences calculated from SAFEP

Where *a* and *b* are generic lipid species, M is a protein conformation, 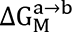 is the free energy cost of transforming a lipid of species *a* to species *b* bound to M, 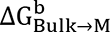 is the free energy cost of transferring a lipid of species *b* from the bulk to a protein in state M, and 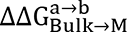 is the relative free energy of exchanging a bound lipid of species *a* with a lipid of species *b* from the bulk.

Free energies of transformation of bound ligands 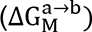 were calculated for each of two conformations (WT and ELIC5) and each of three *a-b* pairs (POPC vs POPE, POPG vs POPE, POPC vs POPG). In this case, the bound pose could be defined using a relaxed conformation of the initial protein-lipid complex. The DBC only included the glycerol atoms of the lipid since inclusion of the more flexible acyl chains and headgroups would require a much looser restraint, thus reducing the sampling benefits.

Following equilibration of the membrane-embedded proteins, the bound lipid was replaced by one of three hybrid (or “dual”) topologies: alchemical lipids which share acyl chains and glycerol cores, but bear two separate phospholipid headgroups. Protein-lipid FEP calculations were carried out with half flat-bottom DBC restraints on the bound lipid to prevent unbinding. The upper wall was placed near the 95^th^ percentile of the unbiased simulation of POPC bound to ELIC5 (the most weakly bound protein-lipid complex), which corresponded to 6 Å (Supplementary Figure 1). 120 lambda windows of 3 ns per window were run in an embarrassingly parallel arrangement. The systems were equilibrated at each lambda value for at least one nanosecond prior to collection of FEP data. All SAFEP calculations used IDWS for simultaneous calculation of the reverse transformation. Due to the computational cost of these simulations, a single replica was run for each lipid transformation for each conformation (6 individual calculations total).

Membrane-only simulations used to calculate 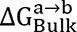 lacked DBC restraints but required two modifications to prevent lipid flipping during FEP. The electrostatic scaling was started at lambda = 0.25 instead of 0.5; this setting prevented complete discharging of the lipid. Forty linearly spaced 1.4 ns FEP windows were used, with 0.5 ns of equilibration prior to collection of FEP data. This modification was not sufficient to prevent lipid flipping for some transformations involving POPE and POPG (Supplementary Figures 3 and 4), and in these cases a flat-bottom harmonic restraint was placed between the central carbon on the FEP lipid and a lipid in the opposite leaflet (lower bound 28.5 Å; upper bound 41.5 Å; force constant 4.0 kcal/mol/Å^2^). Five replicas were run for each lipid comparison in each membrane composition. Replicas were generated by selecting a different lipid for each calculation.

### SAFEP Analysis and Convergence-Testing

The output from the FEP calculations was used to estimate the difference in free energy between the two end states using an accumulated Bennett’s Acceptance Ratio estimator (BAR) as implemented in PyMBAR^67^ using the alchemlyb interface^68^, and custom scripts. Decorrelated samples were obtained using subsampling utilities available in those libraries. Replicate data were aggregated for final free energy estimations. Errors were estimated by using the BAR estimator as implemented in PyMBAR in the case of the protein FEP calculations and by the standard error of the mean of replicas in the case of the membrane FEP calculations. All analysis scripts are available on GitHub^24^.

We considered convergence of the calculations through several metrics. The cumulative change in free energy with respect to λ are shown in Supplementary Figures 3 and 7. We further tested convergence by comparing the cumulative ΔG values that resulted from using the first vs second half of the raw data, for each replica (Supplementary Figures 4 and 8), and required that these values be within ∼1 kcal/mol of each other. Finally, internal hysteresis was assessed by calculating the difference *δ*_λ_ between the forward and backward exponential free energy estimates per window. Well-converged calculations are expected to have only small differences between forward and backward calculations (Supplementary Figures 5 and 9) that are randomly distributed with respect to lambda (Supplementary Figures 6 and 10). We used approximately minimal lambda schedules to ensure convergence thus defined; shorter, fewer windows were found to yield inadequate sampling of the ensembles.

### Pharmacological and allosteric predictions

The relative free energies of replacement calculated above (Equation 1a, Table 2) can then be converted to relative binding constants:

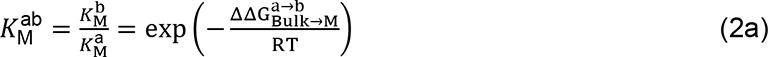

Where 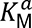 is the absolute binding constant of lipid *a* to state M, and 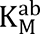 is the relative binding constant of lipid *b* relative to that of lipid *a*. For example, the state-dependent binding constant for POPG binding to the ELIC5 conformation (relative to POPC) is:

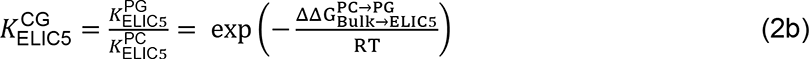

The above can be extended to predictions of allosteric modulation if we make the following assumptions:

1. ELIC has at least two distinct conformations, which we denote ELIC5 and WT.
2. A maximum of three lipid species are present: POPE, POPC, and POPG. The sum of mole fractions of the three lipids is 1: *x*_POPC_ + *x*_POPE_ + *x*_POPG_ = 1
3. The allosteric effects of each binding site are independent of the other binding sites.
4. The probability of a site being unoccupied is negligible (i.e. when a lipid leaves the binding site it is rapidly replaced by another)

Under these assumptions, the probability 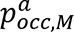 of a site in conformation M to be occupied by a lipid of species *a* is

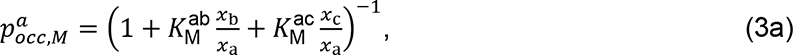

For example, the fraction of sites occupied by POPC in the ELIC5 conformation is

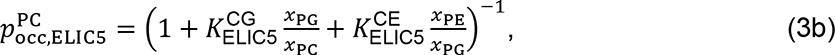

The relative probability of observing the protein in each of the two conformations, as a function of lipid composition, is derived below. First, the law of mass action for a lipid binding to a given conformation is given by the product of the total amount of protein in that state, the equilibrium constant of binding, and the mole fraction of the lipid in the membrane. For example:

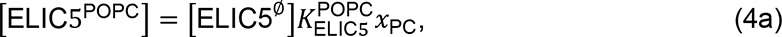

where [ELIC5^POPC^] and [ELIC5^∅^] are the concentrations of ELIC in the ELIC5 state, with bound POPC and without a bound lipid, respectively. The probability ratio of observing ELIC in the

ELIC5 conformation vs the WT conformation 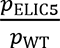 is then:

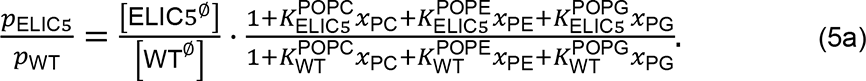

We assume that the fraction of receptors with unoccupied sites is negligible, and switch to POPC-relative binding constants to obtain:

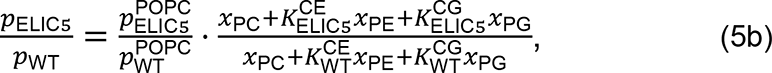

This expression yields the ratio between the overall population of both conformations as the ratio obtained in pure POPC, multiplied by the modulation factor due to the presence of POPE and POPG lipids. We separate the latter contribution and take its logarithm, yielding the quantity plotted in Figure 5C:

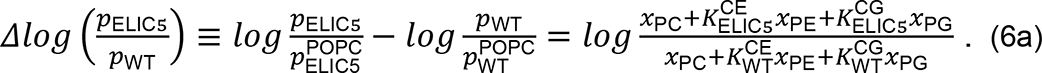

## Acknowledgements

This study was supported by grants R35GM137957 to WC, F32GM139351 to JP, NSF2152059 to GB and ESM, and K08GM139031 to TJ. We are grateful to Drs. Olaf Andersen and Philipp Schmidpeter for help in establishing the fluorescence stopped-flow flux assay and data analysis, and to Dr. Mark Arcario for useful discussions. The authors also acknowledge the Office of Advanced Research Computing (OARC) at Rutgers, The State University of New Jersey for providing access to the Caliburn cluster and associated research computing resources that have contributed to the results reported here. URL: https://oarc.rutgers.edu.

---

**Extended Data Fig. 1.**
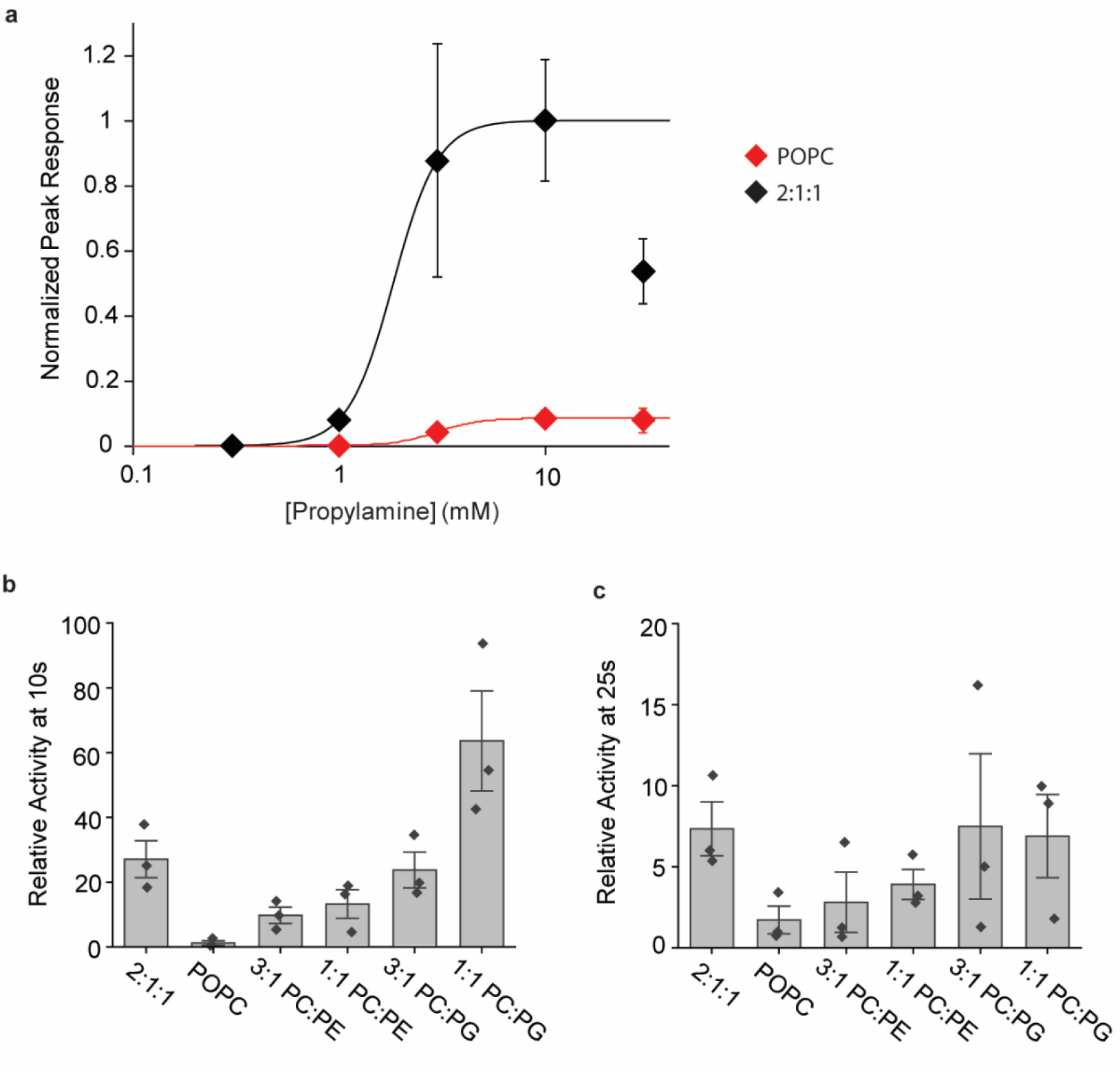
ELIC activity in different lipid conditions. **(a)** Normalized plots of peak rates of ELIC in response to propylamine in 2:1:1 or POPC liposomes in the Tl^+^ flux assay (n = 3, ± SEM). Data are fit to a Hill equation yielding an EC_50_ of 2.4 ± 0.4 mM and n of 2.2 ± 0.7 for 2:1:1, and an EC_50_ of 4.3 ± 1.4 mM and n of 3.6 ± 0.6 for POPC (± SEM). **(b)** Relative activity at 10 s from Tl^+^ flux data in Fig. 1c, measured as the rate at 10 s divided by the peak rate (n = 3, ± SEM). **(c)** Relative activity at 25 s from Tl^+^ flux data in Fig. 1c, measured as the rate at 25 s divided by the peak rate (n = 3, ± SEM).

**Extended Data Fig. 2.**
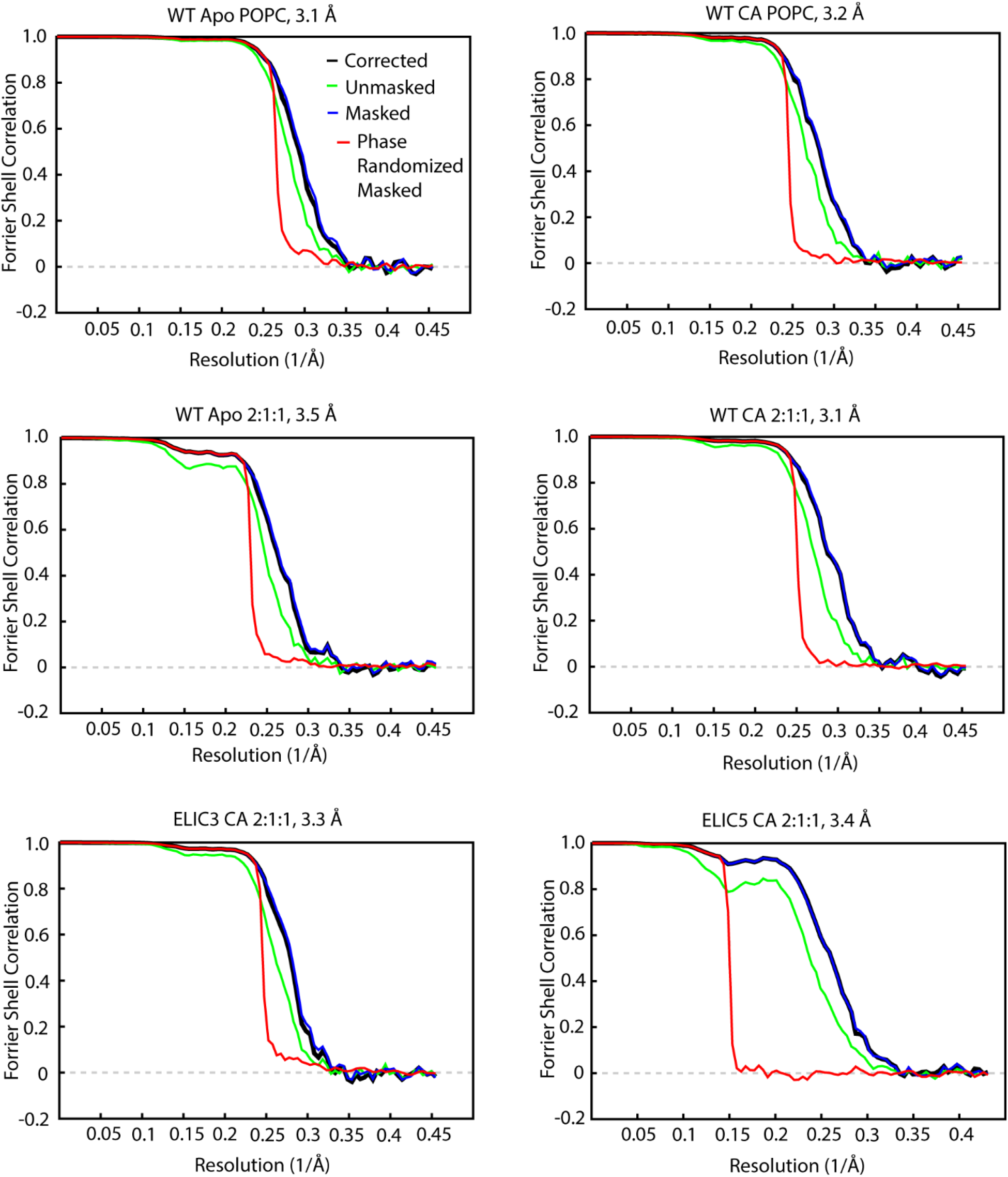
Fourier shell correlation curves. FSC curves for each ELIC structure labeled according to WT ELIC, ELIC3, ELIC5; apo or with agonist (10 mM cysteamine, CA); the lipid used in the nanodisc (POPC or 2:1:1); and the final post-processed resolution.

**Extended Data Fig. 3.**
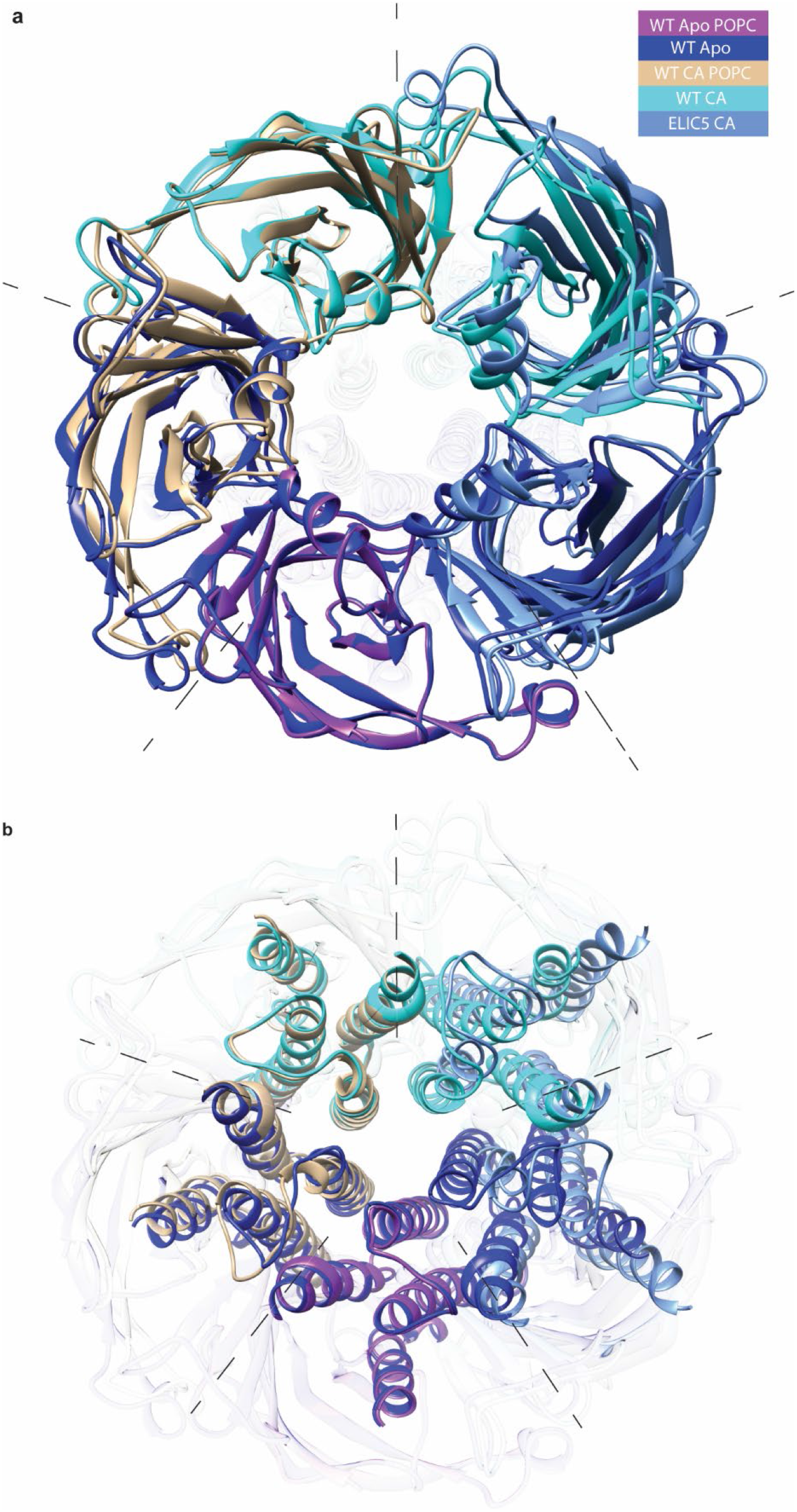
ELIC structural comparisons. **(a)** Overlapped ECD of indicated pairs of ELIC structures, including WT apo POPC (WT apo in POPC nanodiscs, WT apo (WT apo in 2:1:1 nanodiscs), WT CA POPC (WT + 10 mM cysteamine in POPC nanodiscs), WT CA (WT + 10 mM cysteamine in 2:1:1 nanodiscs), ELIC5 CA (ELIC5 + 10 mM cysteamine in 2:1:1 nanodiscs) **(b)** Overlapped TMD of indicated pairs of ELIC structures.

**Extended Data Fig. 4.**
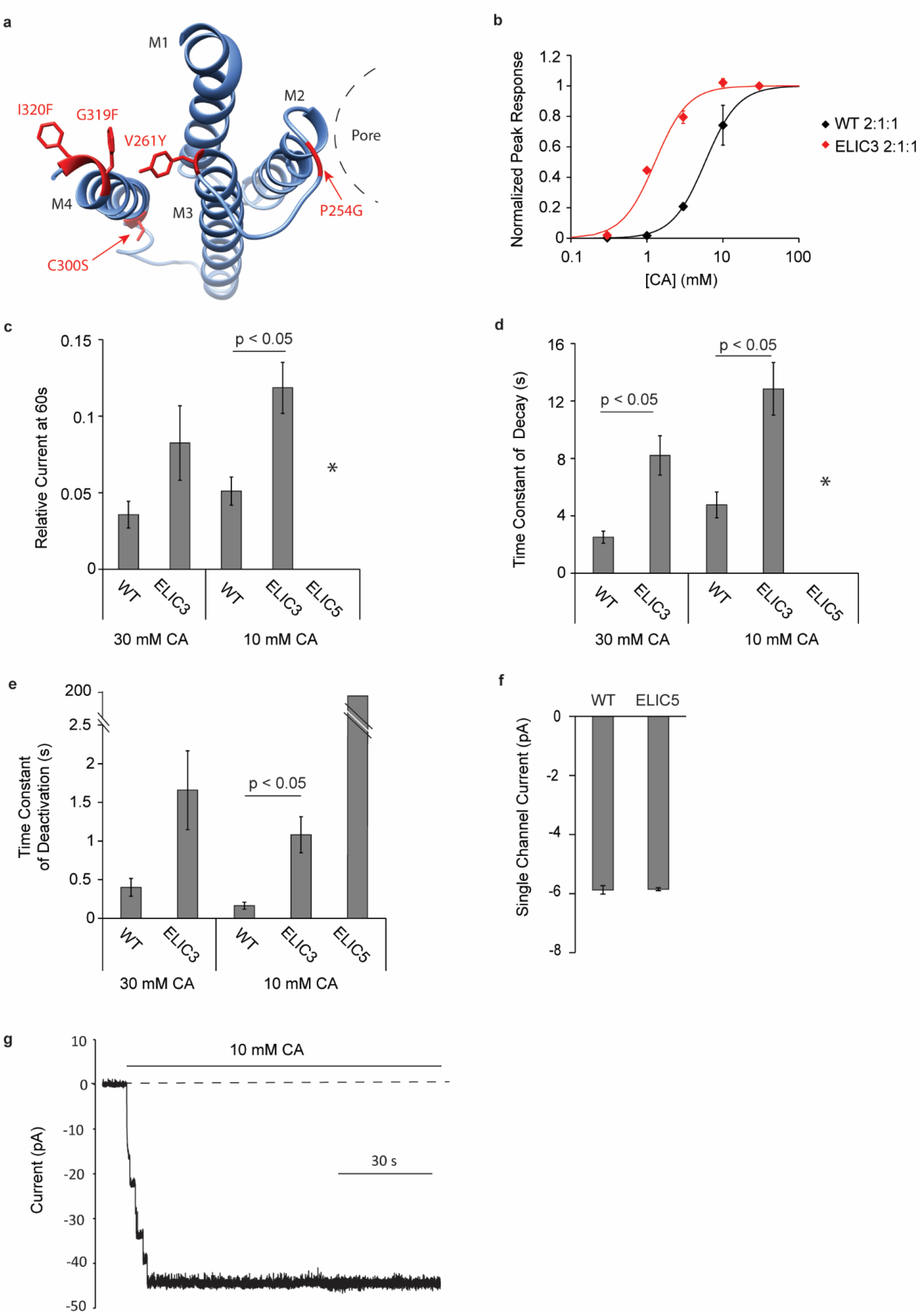
Functional characterization of ELIC gain-of-function mutants. **(a)** View of one subunit of ELIC TMD from the extracellular side showing the mutated residues (red) for ELIC3 (V261Y, G319F, I320F) and ELIC5 (V261Y, G319F, I320F, P254G and C300S). **(b)** Peak dose response curve of WT ELIC and ELIC3 with cysteamine (CA) obtained from excised patch-clamp recordings from 2:1:1 giant liposomes. Data are fit to a Hill equation with n set to 2, yielding an EC_50_ of 5.2 ± 0.3 mM for WT ELIC and 1.2 ± 0.2 mM for ELIC3 (n = 4, ± SEM). Of note, the recording buffer contained 0.5 mM BaCl_2_, to maintain patch stability, which leads to ∼2-3x right-shift in the EC ^70^. **(c)** Relative current at 60 s (current at 60 s divided by the peak current) after the application of 10 or 30 mM cysteamine from giant liposome recordings of WT ELIC and ELIC3 (n = 4, ± SEM). * indicates no data because there was no evidence of current decay in ELIC5 responses. **(d)** Same as (c) showing the weighted time constant of decay in WT ELIC and ELIC3 currents in response to 10 or 30 mM cysteamine (n = 4, ± SEM). **(e)** Time constant of deactivation after 1 s application of 10 or 30 mM cysteamine in WT ELIC, and ELIC3 giant liposome recordings. For ELIC5, 10 mM cysteamine was applied for ∼1 min, achieving stable current, prior to removal of agonist (n = 4-11, ± SEM). **(f)** Single channel currents from WT and ELIC5 giant liposome recordings at −60 mV (n = 4, ± SEM). **(g)** Representative ELIC5 response to 10 mM cysteamine from a giant liposome recording, in which single channel openings were resolved. No closing events were observed for ∼2 min.

**Extended Data Fig. 5.**
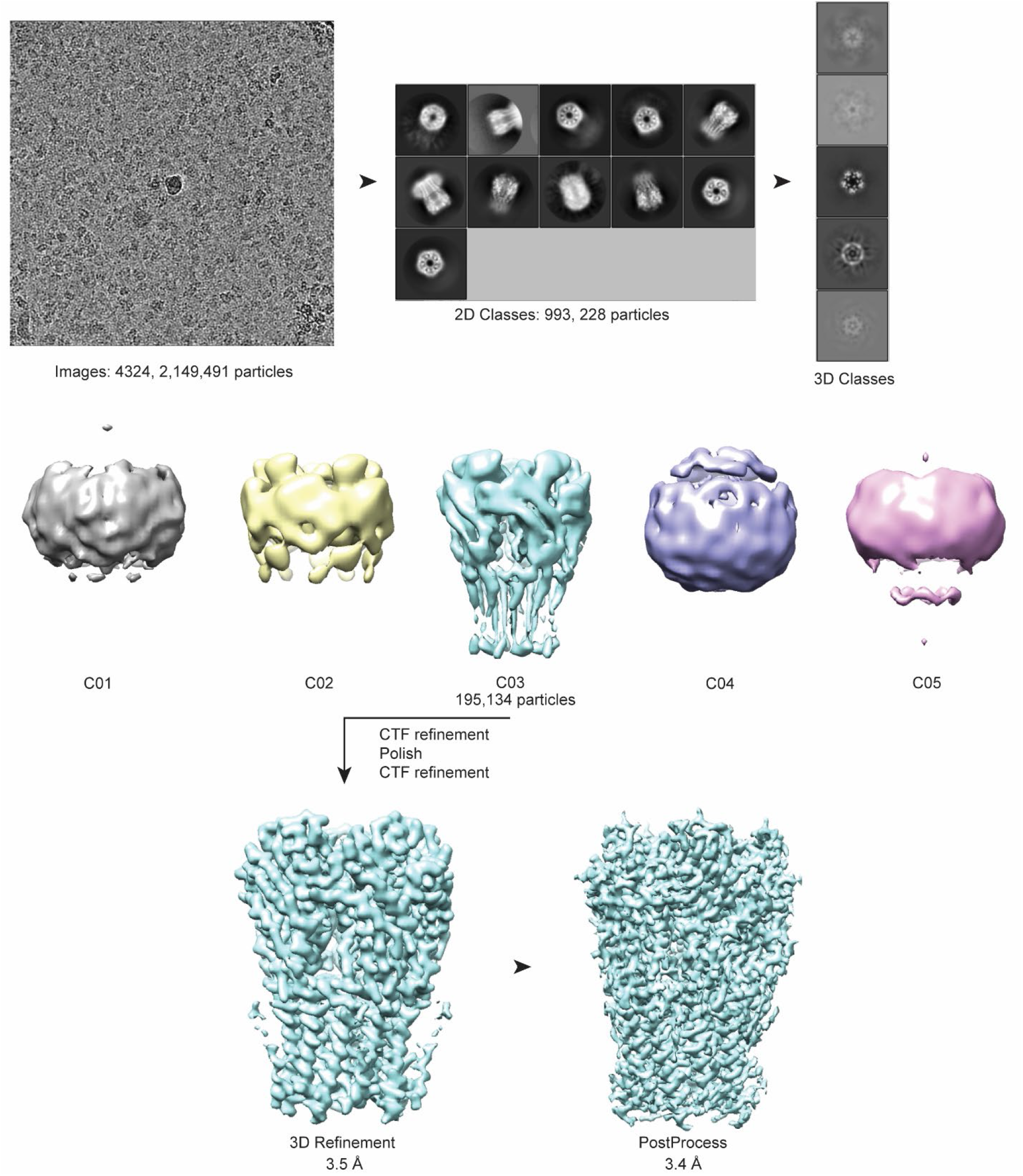
Diagram for single particle analysis of the ELIC5 CA structure using Relion3. Workflow for analysis showing a representative micrograph, 2D classes, 3D classes and refined and post-processed maps. This workflow for the ELIC5 CA structure is representative of the process for all other structures.

**Extended Data Fig. 6.**
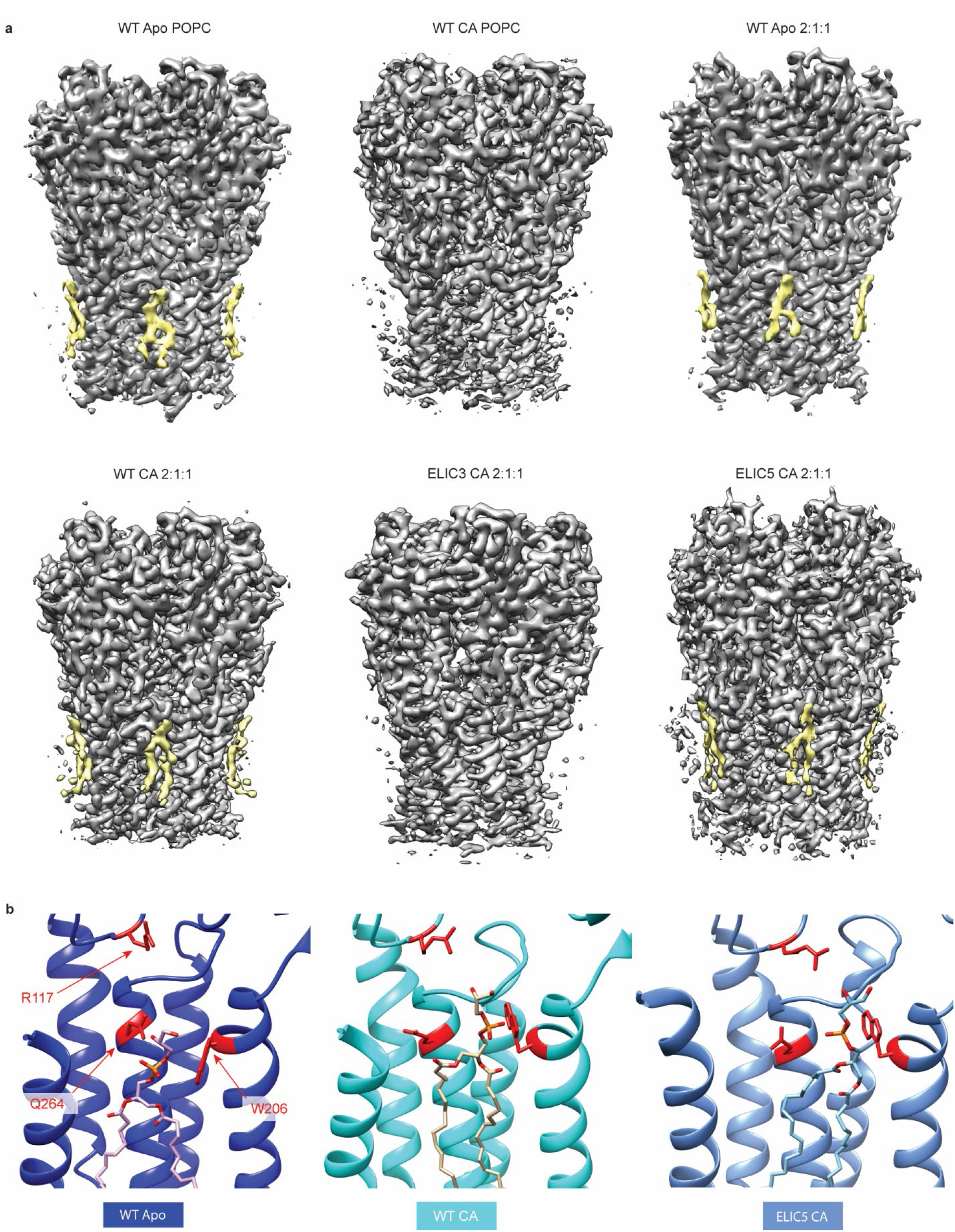
Bound phospholipids in ELIC structures. **(a)** Electron density maps of all ELIC structures with and without agonist (10 mM cysteamine) in POPC or 2:1:1 nanodiscs. Shown in yellow is the density for a bound phospholipid illustrating its location at the outer leaflet site. The threshold of the electron density was adjusted to optimize the appearance of the phospholipid density, and thus the strength of the phospholipid density is not comparable between structures. No significant phospholipid-like densities were detected at other sites. **(b)** Binding modes of the phospholipid in the indicated structures highlighting the position of R117, Q264 and W206 relative to the phospholipid in each structure. Q264 and W206 closely interact with the phospholipid headgroup in the WT apo structure and rotate away from the lipid-protein interface in the WT CA and ELIC5 CA structures. R117 is closest to the phospholipid headgroup in the ELIC5 CA structure.

**Extended Data Fig. 7.**
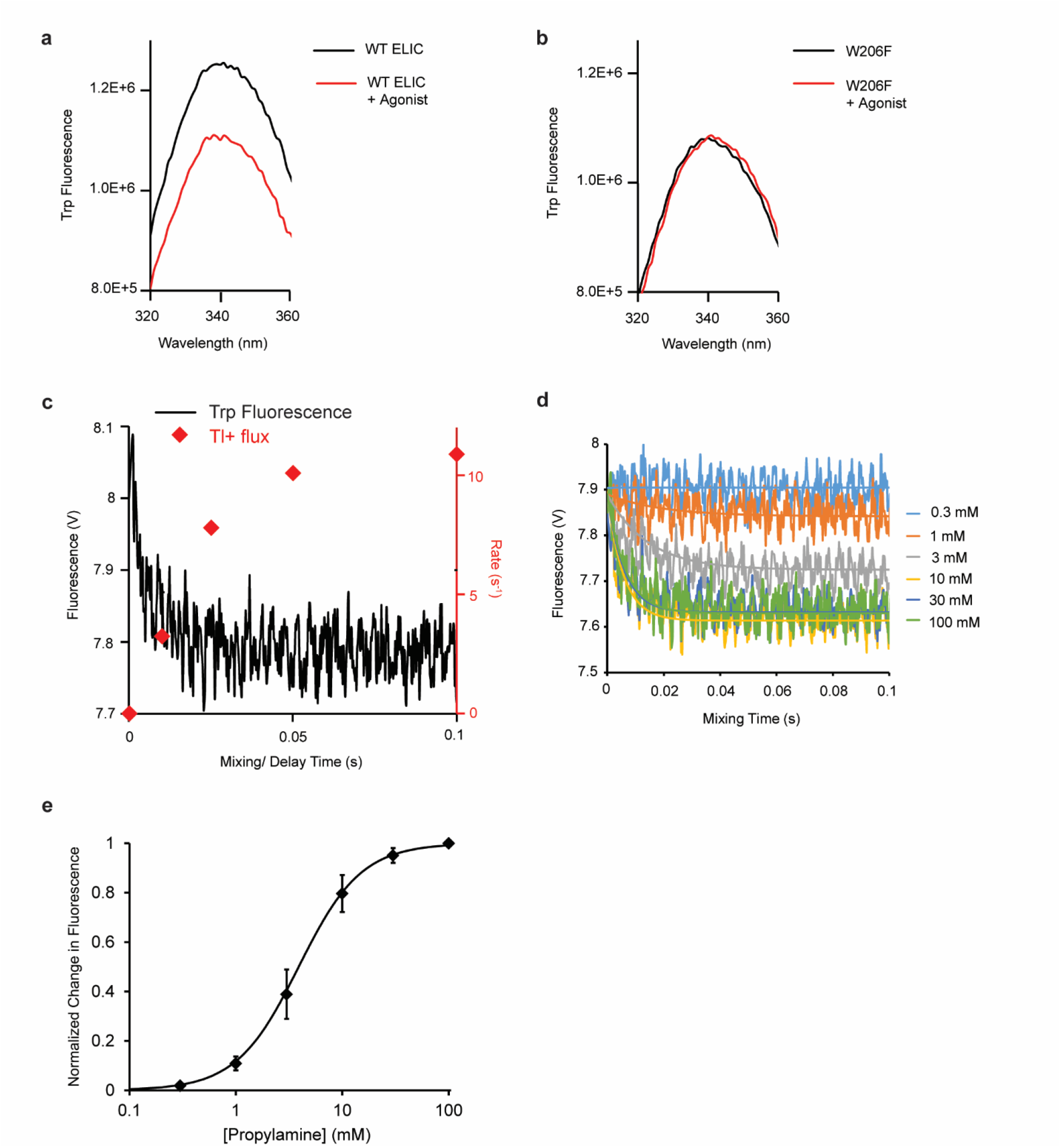
Stopped-flow tryptophan (TRP) fluorescence measurements of ELIC. **(a)** Emission spectrum of WT ELIC with and without 10 mM propylamine (excitation 295 nm). **(b)** Same as (a) with W206F ELIC. **(c)** Time course of TRP fluorescence of WT ELIC after rapid mixing with 10 mM propylamine (black, time constant = 9.9 ± 0.6 ms, n = 3, ± SEM). The sample was excited at 295 nm to monitor TRP fluorescence. Also shown is the time course of WT ELIC activation by Tl^+^ flux in response to 10 mM propylamine in 2:1:1 liposomes (red, data obtained from Fig. 1, time constant = 41 ± 34 ms, n = 3, ± SEM). In POPC liposomes, the time constant for WT ELIC activation by Tl^+^ flux in response to 10 mM propylamine was 20 ± 6 ms (n = 3, ± SEM). **(d)** Representative time courses of TRP fluorescence of WT ELIC after rapid mixing with varying concentrations of propylamine. **(e)** Normalized change in TRP fluorescence derived from time courses as shown in (d). Data are fit to a Hill equation yielding EC_50_ = 4.3 ± 0.8 mM and n = 1.6 ± 0.3 (n = 3, ± SEM). These data indicate that movement of W206 shows a similar dependence on propylamine concentration as channel opening (Extended Data Fig. 1).

**Extended Data Fig. 8.**
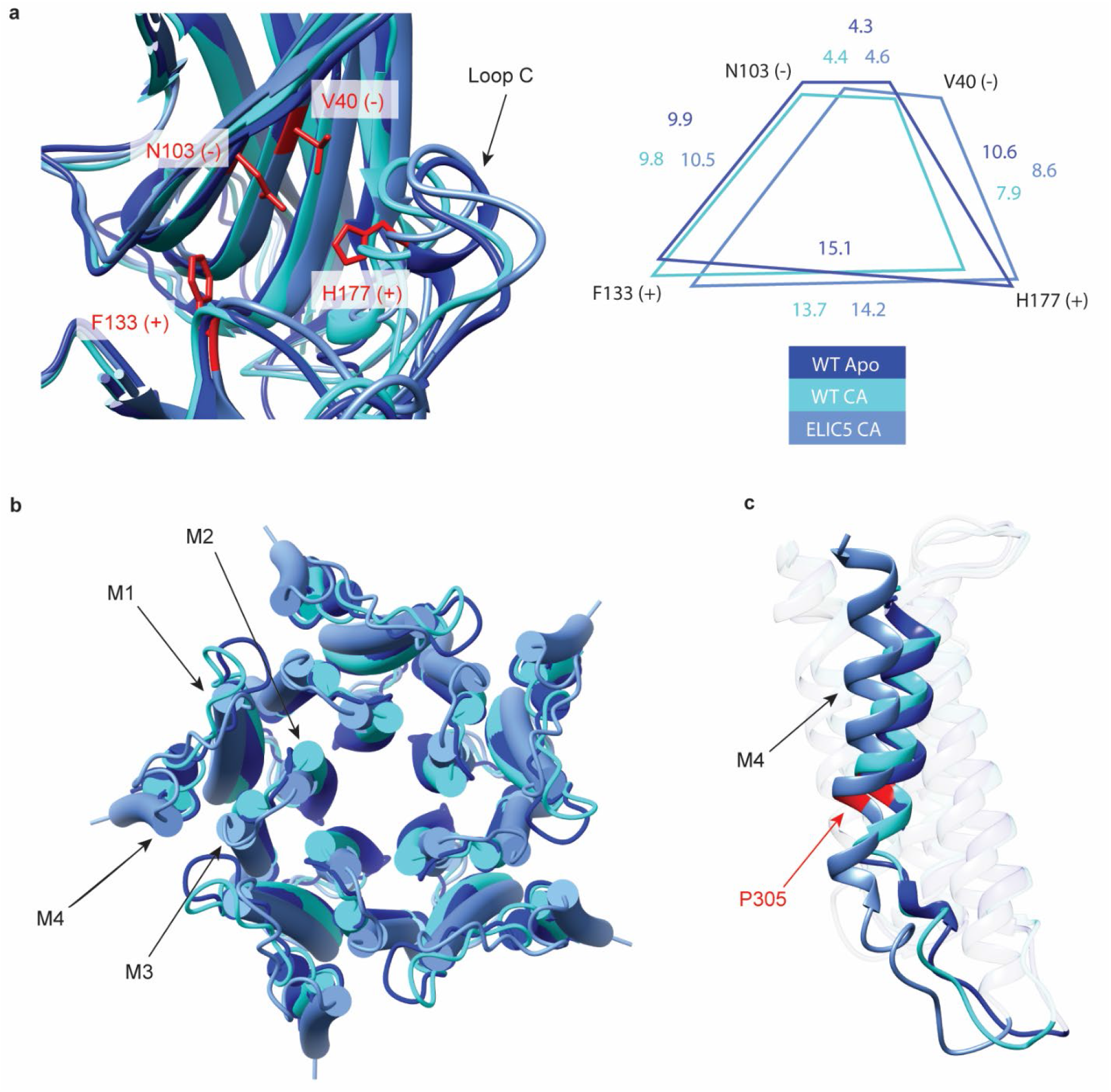
Comparison of the agonist binding site and TMD between ELIC structures. **(a)** *Left:* Representation of the agonist binding site in WT apo, WT CA and ELIC5 CA structures highlighting the conformational change in loop C. Shown in red are four key residues that form the agonist binding site. *Right*: Distances (Å) between four key residues within the agonist binding site showing contraction of the site in the agonist-bound structures compared to WT apo, and outward translation of the site in the ELIC5 CA structure compared to WT CA. **(b)** View of the TMD from the intracellular side of WT apo, WT CA and ELIC5 CA structures. **(c)** Comparison of M4 between the indicated structures. Highlighted in red is P305.

**Extended Data Fig. 9.**
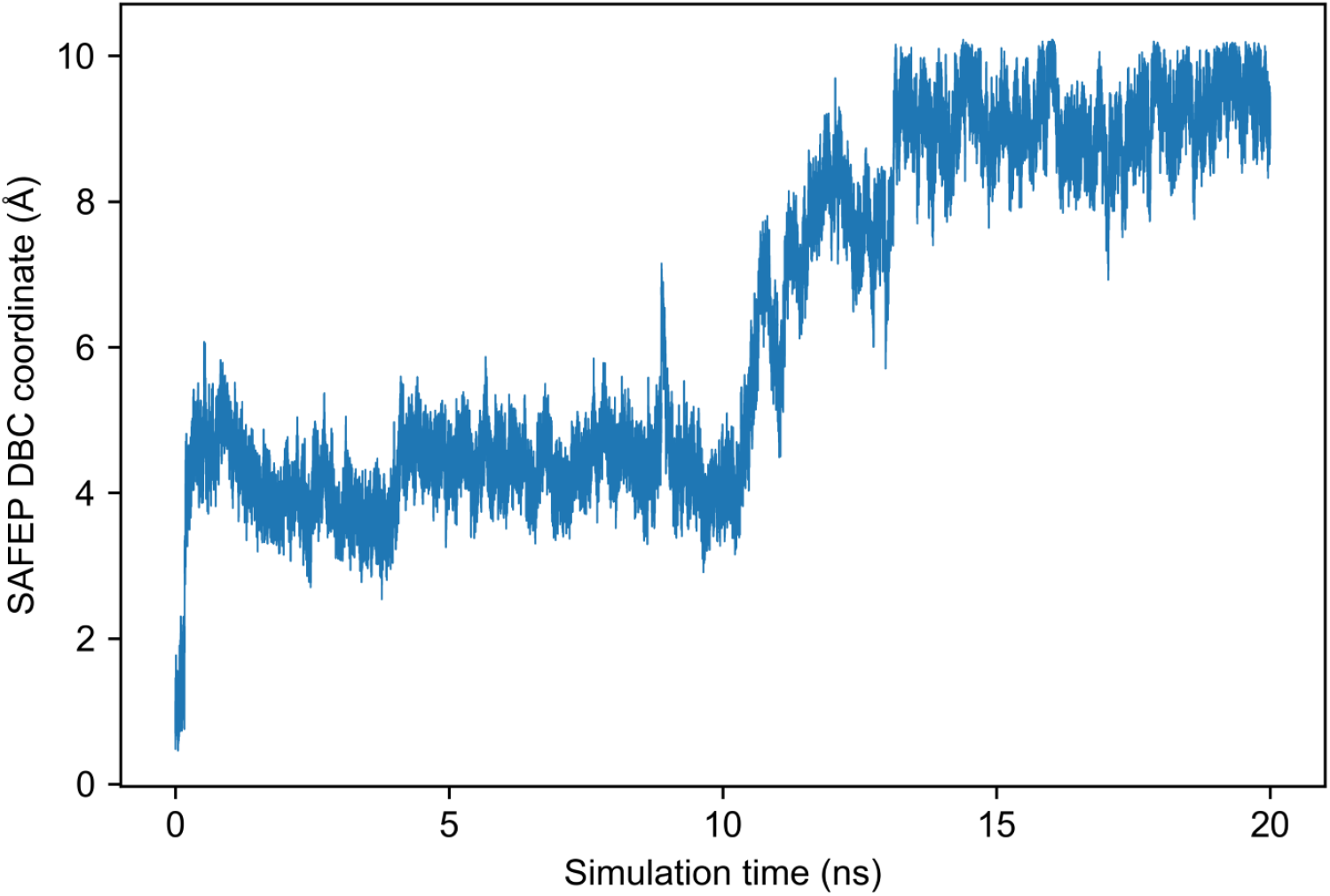
POPG headgroup distance from bound conformation in the WT apo structure. Equilibrium MD simulation demonstrating spontaneous unbinding of the POPG headgroup from the binding mode of the lipid density in the WT apo structure embedded in a 2:1:1 POPC:POPE:POPG membrane, as measured by SAFEP DBC coordinate, which measures the RMSD of selected headgroup glycerol atoms from an initial bound conformation. No bias is applied to POPG until the SAFEP DBC coordinate reaches 10 Å.

**Extended Data Fig. 10.**
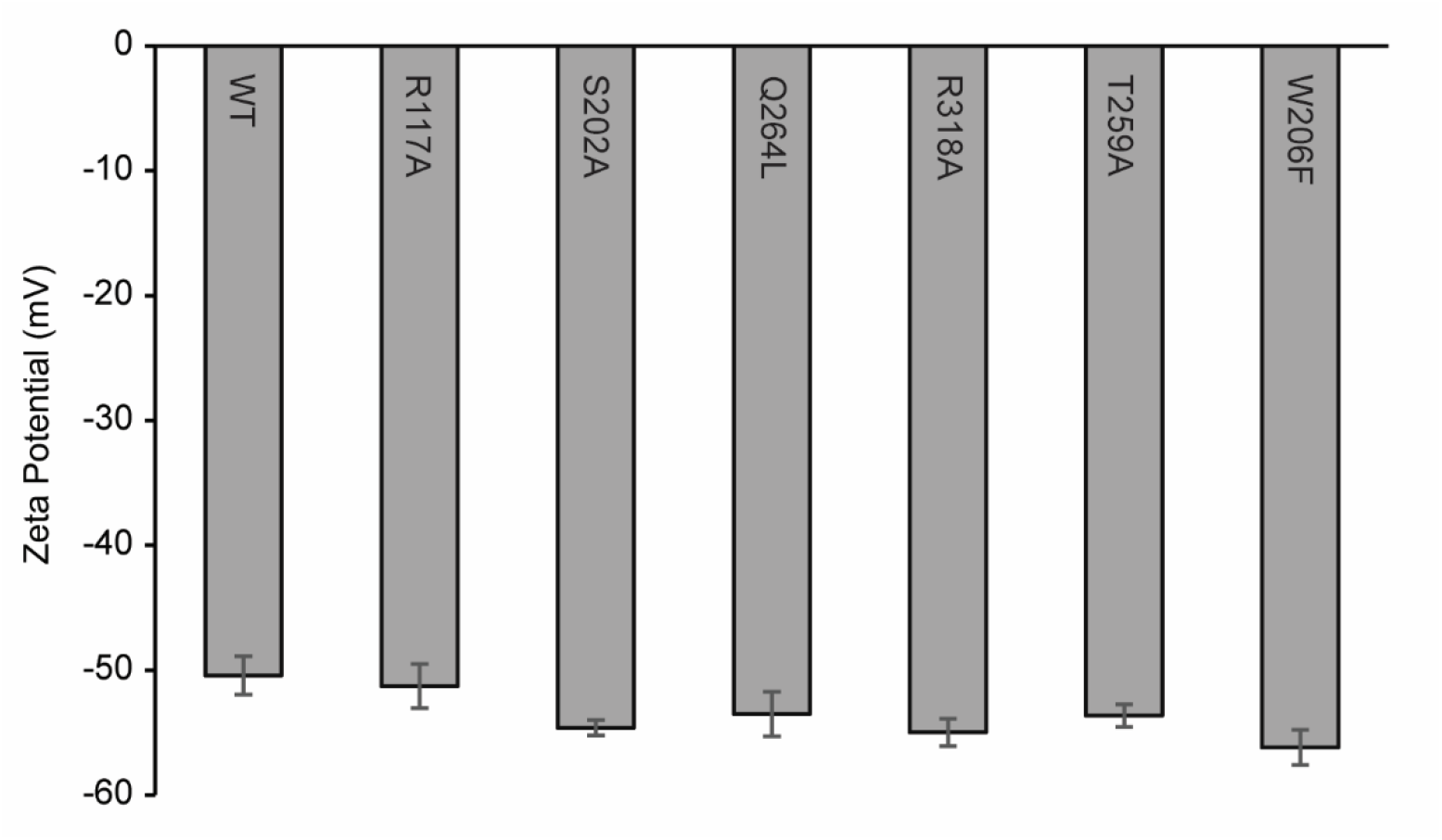
Zeta potential measurements of mutants. Zeta potential values as in Fig. 4b of WT (equivalent to the 3:1 PC:PE Ex ELIC sample) and the indicated mutants. Each sample consists of ELIC in 3:1 POPC:POPE liposomes treated with POPG and methyl-β-cyclodextrin to introduce 25% POPG to the outer leaflet (±SEM). The zeta potential values show that complete POPG exchange was achieved in each of the mutant proteoliposome samples similar to WT.

