## Supplementary Information for "Structural mechanism of leaflet-specific phospholipid modulation of a pentameric ligand-gated ion channel"

**Supplementary Figure 1**

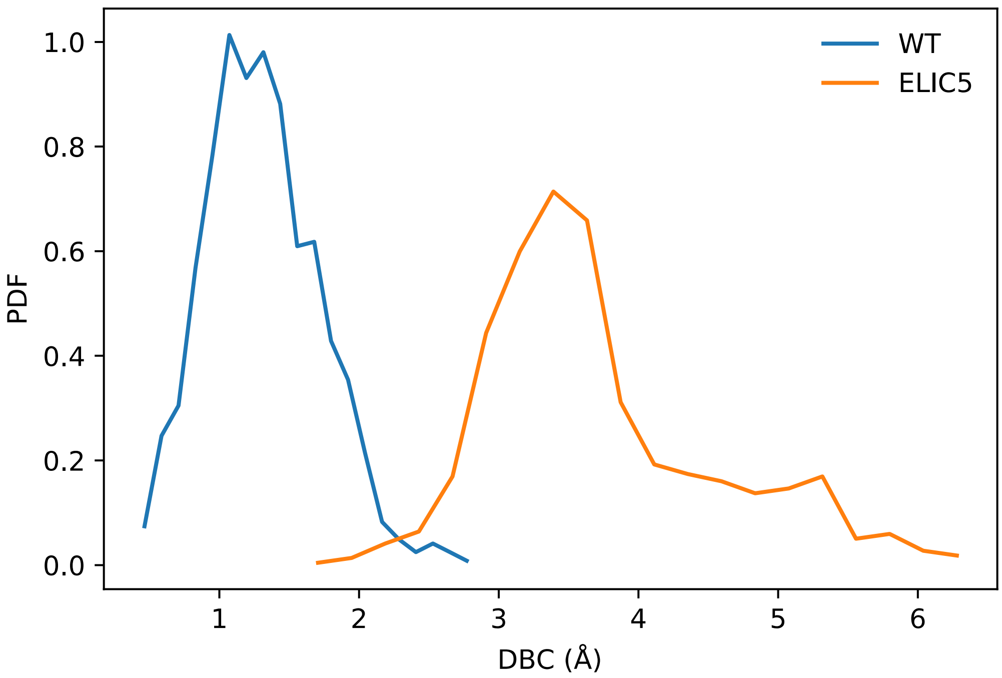


**Supplementary Figure 1 |** Distribution of the DBC collective variable from equilibrium MD simulations of POPC bound to WT and ELIC5. The 95th percentile is approximately 6 Å for ELIC5.

**Supplementary Figure 2**

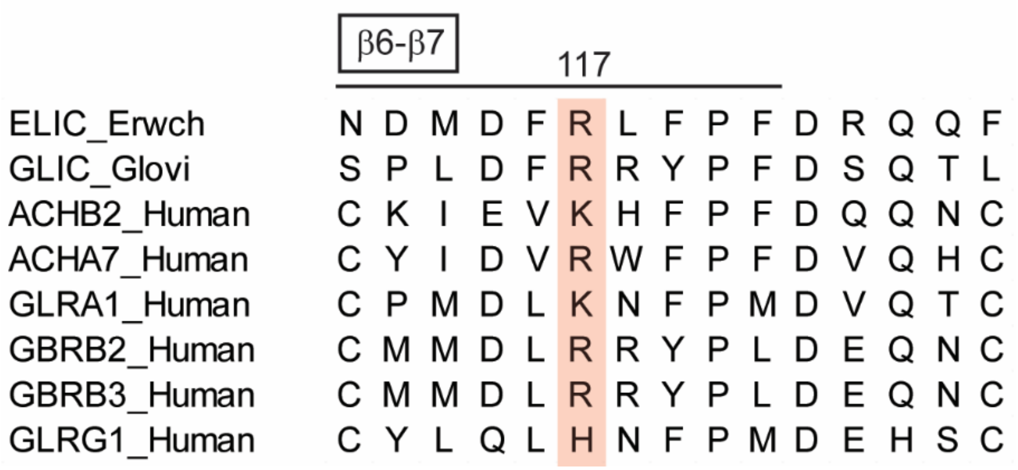


**Supplementary Figure 2 |** Sequence alignment comparing the β6-β7 loop in ELIC with GLIC and other human pLGICs. Highlighted is R117 in ELIC.

**Supplementary Figure 3**

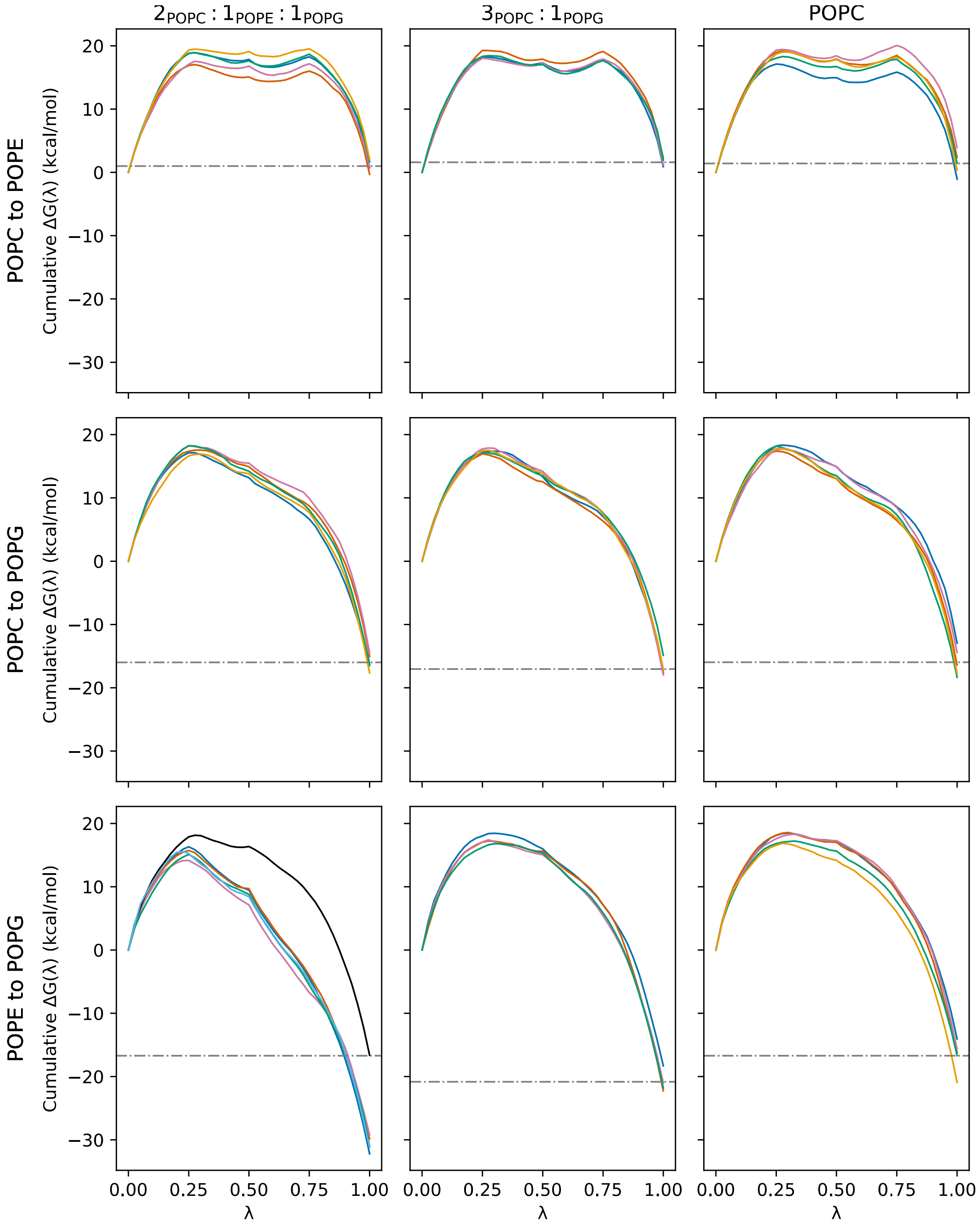


**Supplementary Figure 3 |** Cumulative change in free energy vs λ during SAFEP calculation of $\Delta G_{bulk}$. Calculations are shown for at least 5 replicas of three transformations (rows) in three membrane compositions (columns). Solid lines indicate the cumulative change in free energy with respect to λ for each system, colored by replica. Dashed lines indicate the final average of $\Delta G_{bulk}$ values across replicas, with one exception. The original five replicas of the POPE to POPG transformation in the $2_{POPC}:1_{POPE}:1_{POPG}$ membrane (bottom left) yielded values that were inconsistent with the other two transformations and did not close the cycle. This appears to be due to the tendency of both the PE and PG headgroups to become buried in the hydrophobic core around λ=0.25. We countered this tendency by including a distance restraint between the FEP lipid and another lipid in the opposite leaflet ($2_{POPC}:1_{POPE}:1_{POPG}$, POPE to POPG, black line), yielding the value of $\Delta G_{bulk}$ shown in Table 2 and used in Figure 5.

### Supplementary Figure 4

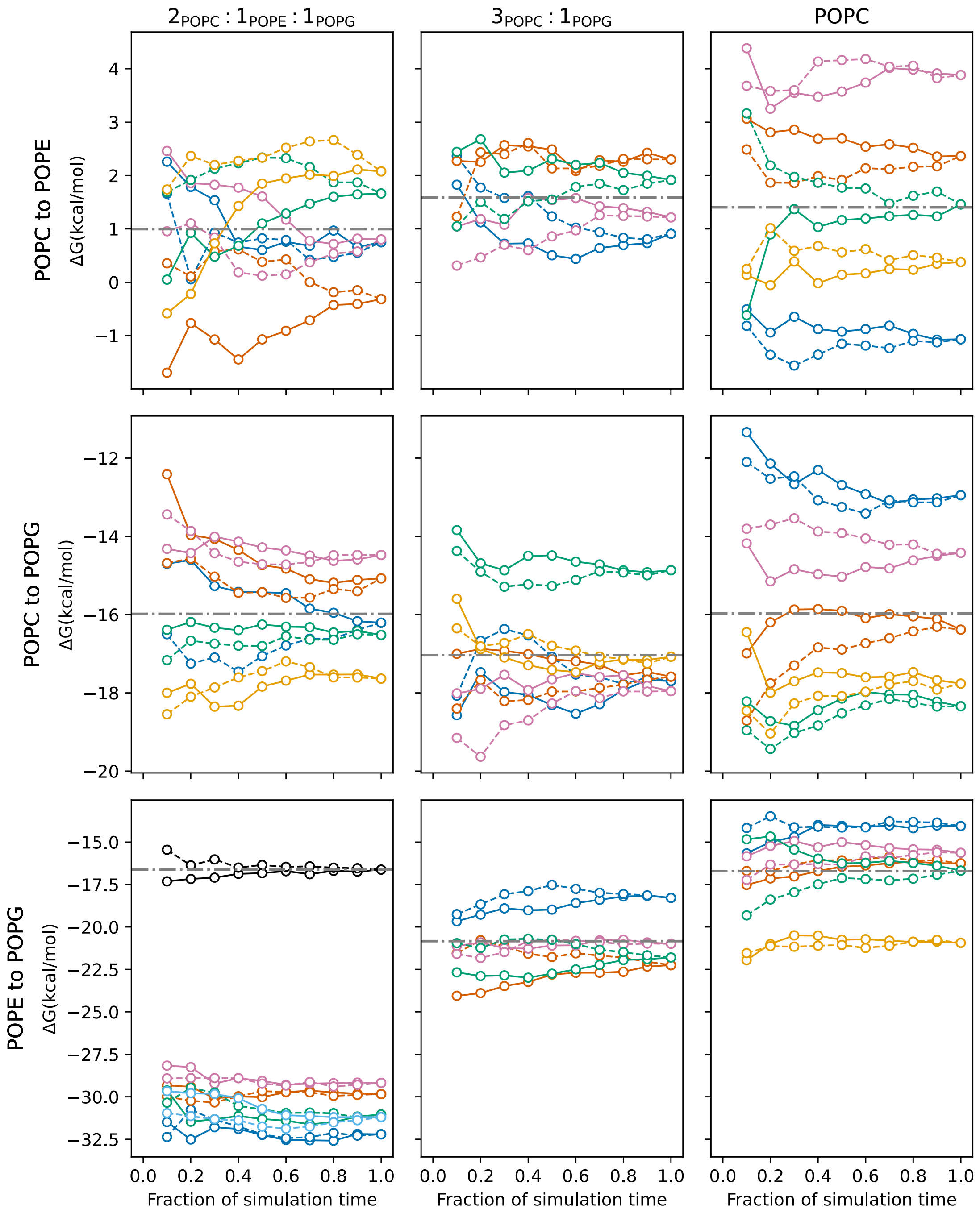


**Supplementary Figure 4 |** Dependence of $\Delta G_{bulk}$ on the fraction of simulation trajectory used for analysis. Dashed lines indicate reverse-time subsampling (starting from the end of the simulation) while solid lines indicate forward-time subsampling (starting from the end of equilibration for all windows). Subsamples begin to overlap when the fraction of simulation time reaches 0.5. Colors indicate individual replicas. Dot-dashed lines indicate final average values across replicas, with one exception. The original five replicas of the POPE to POPG transformation in the $2_{POPC}:1_{POPE}:1_{POPG}$ membrane (bottom left) yielded values that were inconsistent with the other two transformations and did not close the cycle. This appears to be due to the tendency of the PE headgroup to become buried in the hydrophobic core starting at around $\lambda=0.25$. This can be countered by inclusion of a distance restraint between the FEP lipid and another lipid in the opposite leaflet ($2_{POPC}:1_{POPE}:1_{POPG}$, POPE to POPG, black line).

### Supplementary Figure 5

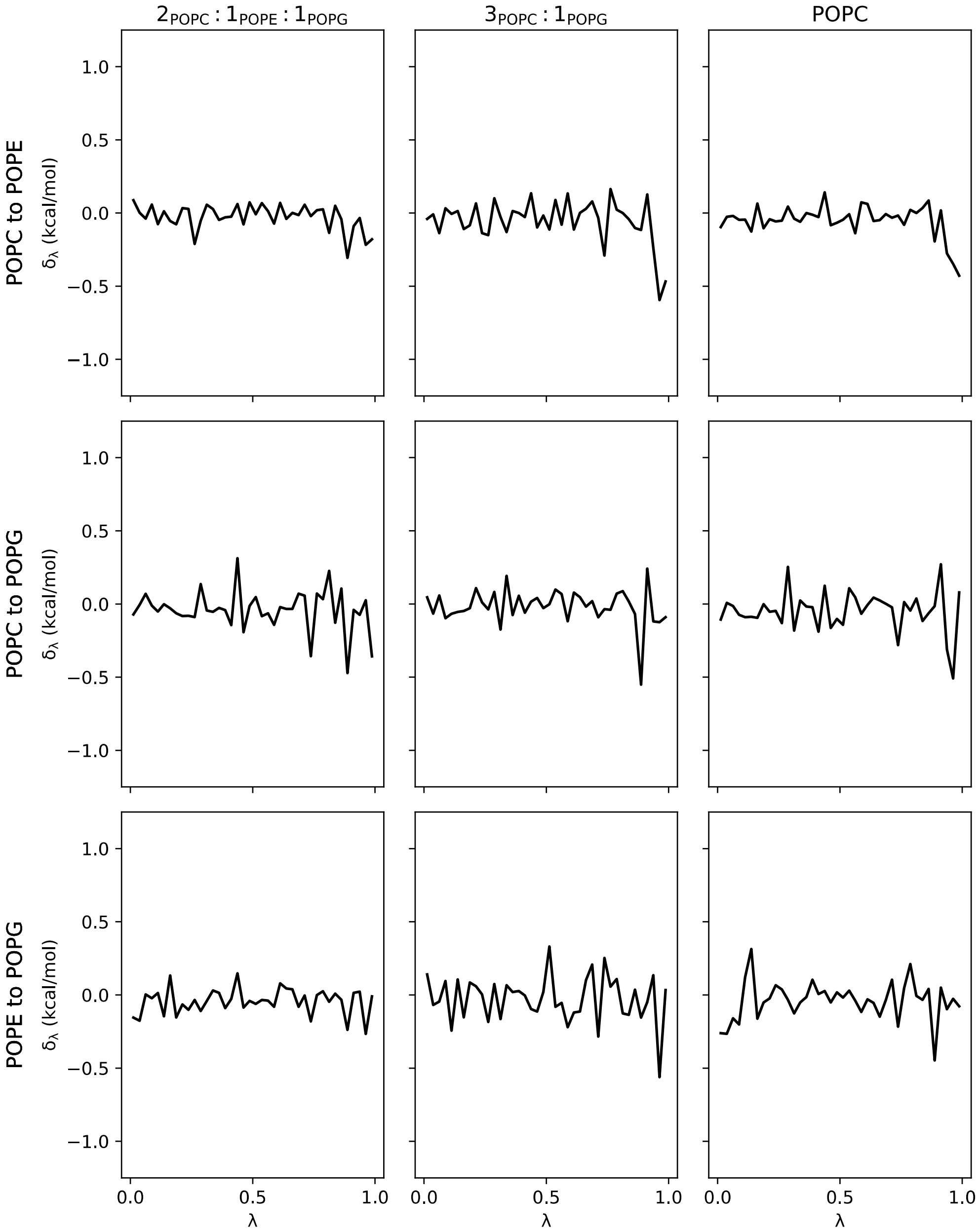


**Supplementary Figure 5 |** Hysteresis in the calculation of $\Delta G_{bulk}$ as a function of λ. Lines indicate the difference ($\delta_\lambda$) between the forward and IDWS-generated backward estimates for each window, using an exponential estimator. Colors represent individual replicas; black curves are determined using all replicas. $\delta_\lambda$ values appear to be independent of λ, suggesting the systems are well-equilibrated with respect to λ.

**Supplementary Figure 6**

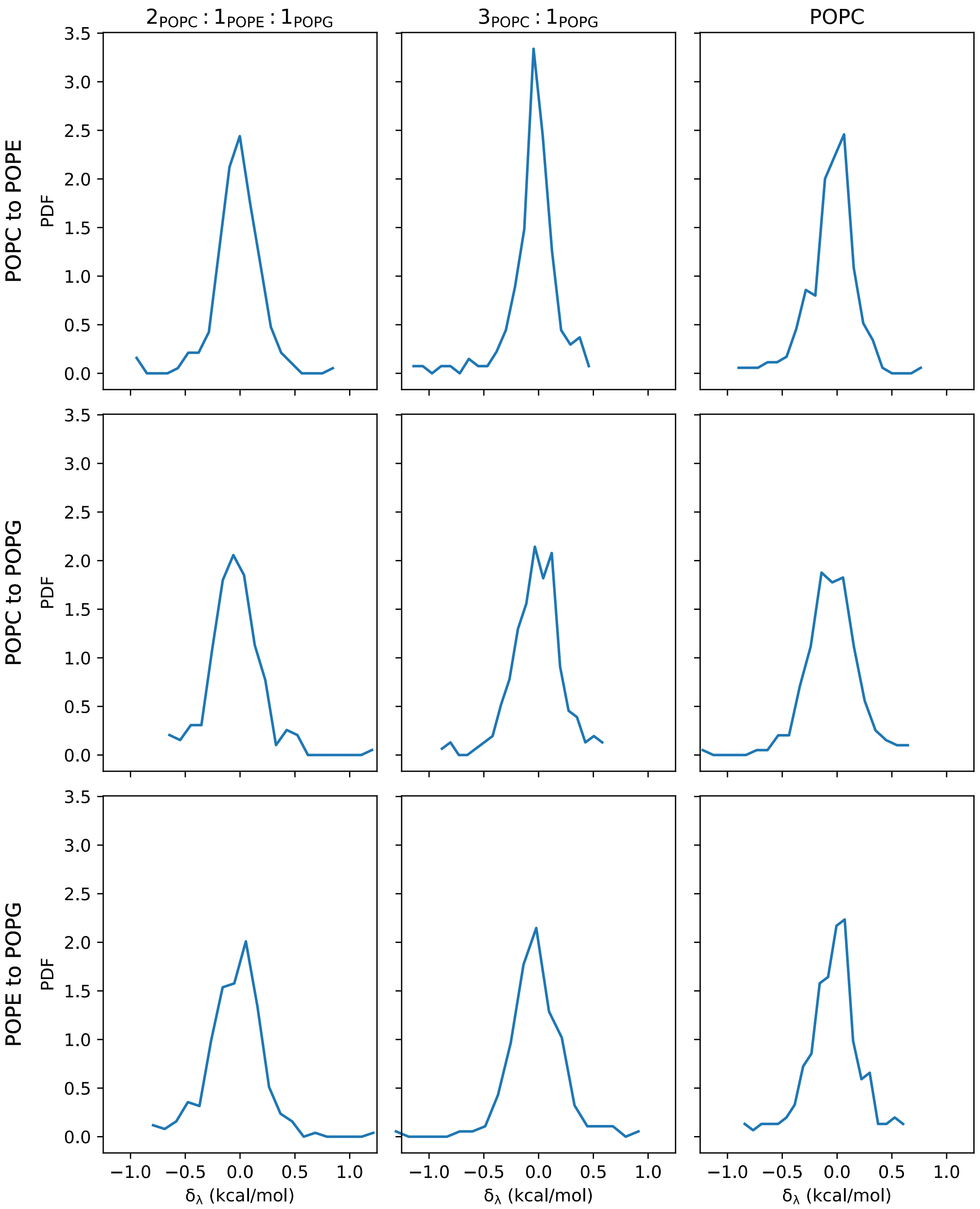


**Supplementary Figure 6 |** Distributions of aggregate $\delta_\lambda$ values shown in Supplementary Figure 5. Values are roughly centered around 0, indicating that the aggregate calculations are well-sampled.

**Supplementary Figure 7**

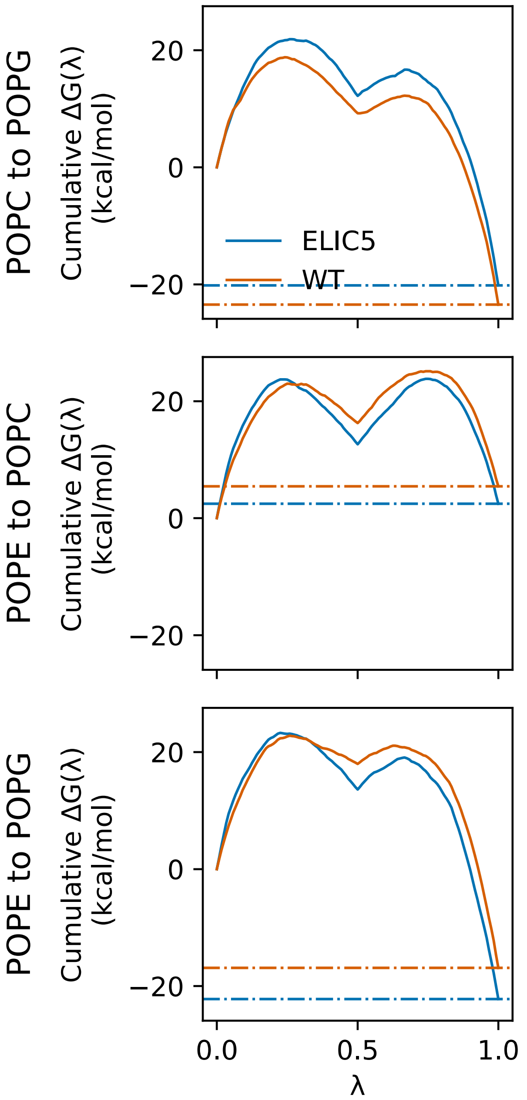


**Supplementary Figure 7 |** Cumulative change in free energy vs λ during SAFEP calculation of $\Delta G_M$. Calculations are shown for three lipid transformations (rows). Solid lines indicate the cumulative change in free energy with respect to λ for each system, colored by protein structure. Dashed lines indicate the final average of $\Delta G_M$ values. As expected, all paths are reasonably smooth; a strong cusp at 0.5 is not unusual because that is the inflection point for both electrostatics and Van der Waals interactions in the protein FEP calculations.

### Supplementary Figure 8

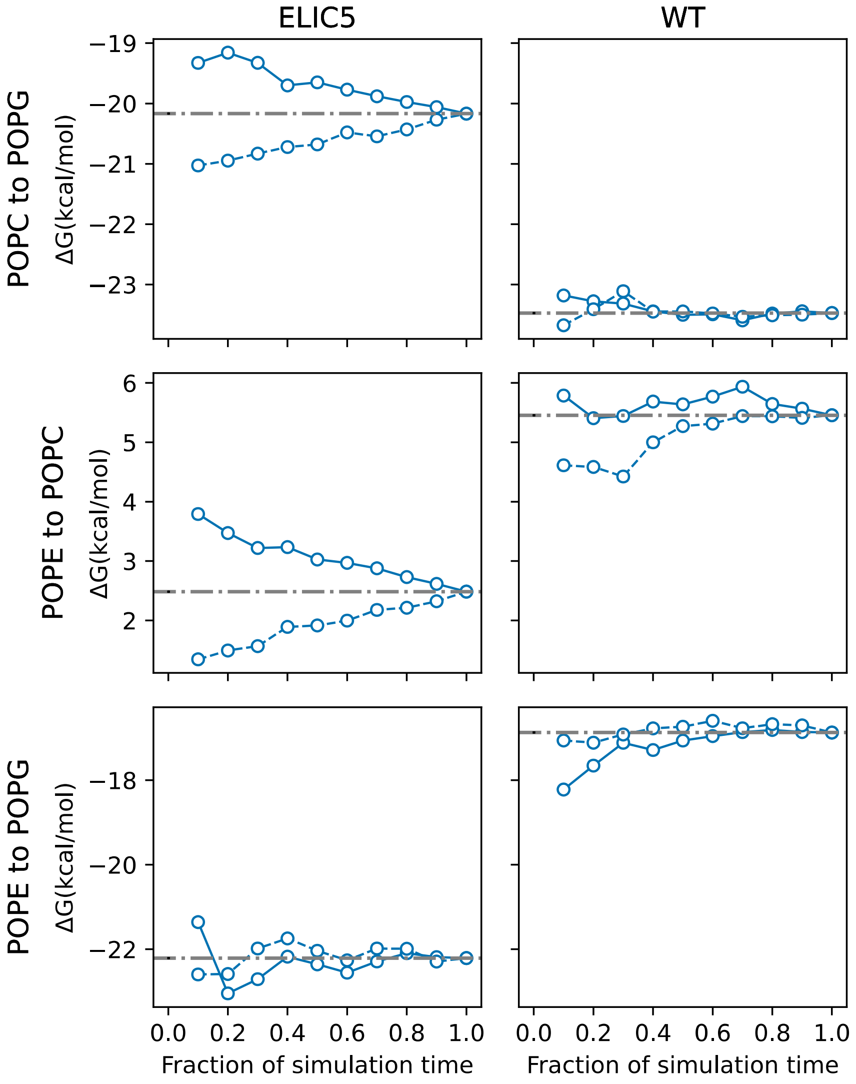


**Supplementary Figure 8 |** Dependence of $\Delta G_M$ on the fraction of simulation trajectory used for analysis, for three lipid transformations (rows) and two protein conformations (columns). Dashed lines indicate reverse-time subsampling (starting from the end of the simulation) while solid lines indicate forward-time subsampling (starting from the end of equilibration for all windows). Subsamples begin to overlap when the fraction of simulation time reaches 0.5. Dot-dashed lines indicate final average values. Well-converged systems are those with a discrepancy of at most 1 kcal/mol at 50% simulation time.

**Supplementary Figure 9**

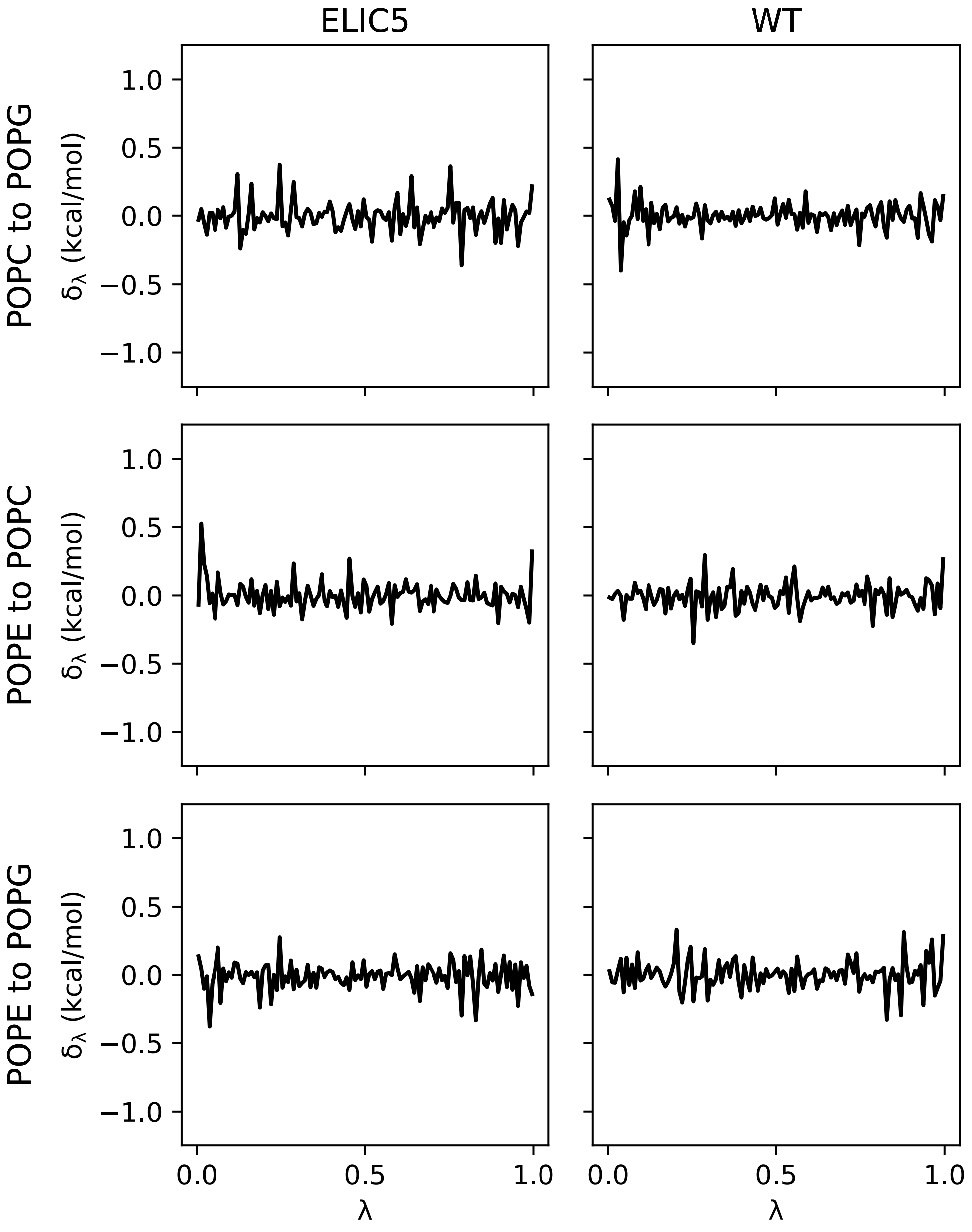


**Supplementary Figure 9 |** Hysteresis in the calculation of $\Delta G_M$ as a function of λ. Lines indicate the difference ($\delta_\lambda$) between the forward and IDWS-generated backward estimates for each window using an exponential estimator. $\delta_\lambda$ values appear to be independent of λ, suggesting the systems are well-equilibrated with respect to λ.

**Supplementary Figure 10**

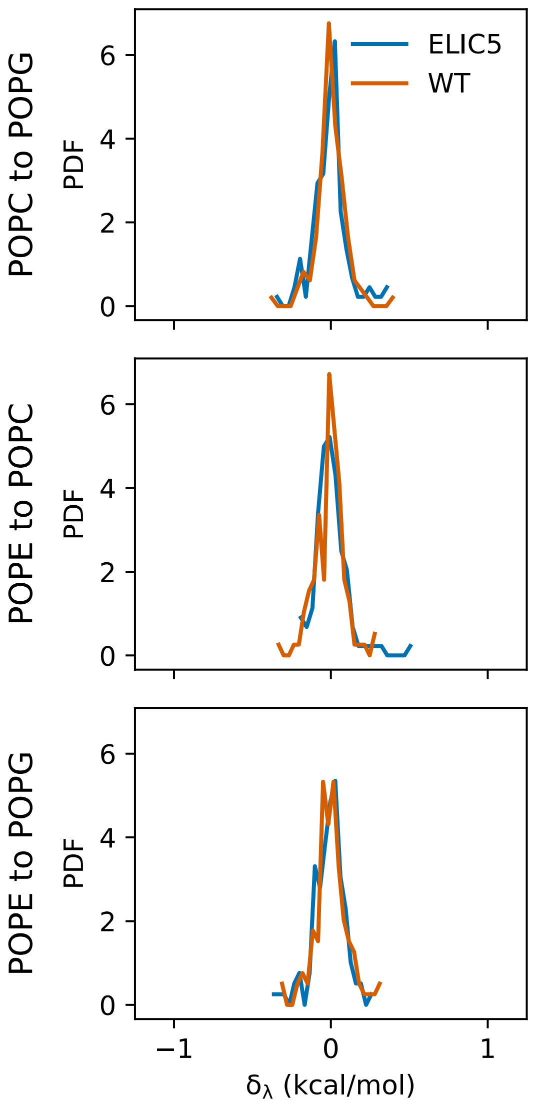


**Supplementary Figure 10 |** Distributions of aggregate protein $\delta_\lambda$ values shown in Supplementary Figure 9. Lines indicate the approximate probability density function for the distribution of each transformation and system (ELIC5 in blue, WT in orange). All $\delta_\lambda$ distributions are symmetric and strongly peaked around 0. The distributions are much narrower than for the membrane calculations, due to the use of significantly more windows (120 vs. 40).
